# Characterization of antimalarial activity of artemisinin-based hybrid drugs

**DOI:** 10.1101/2024.01.26.577447

**Authors:** Helenita Costa Quadros, Lars Herrmann, Jeanne Manaranche, Lucie Paloque, Mariana C. Borges-Silva, Godwin Akpeko Dziwornu, Sarah D’Alessandro, Kelly Chibale, Nicoletta Basilico, Françoise Benoit-Vical, Svetlana B. Tsogoeva, Diogo Rodrigo M. Moreira

## Abstract

In response to the spread of artemisinin (ART) resistance, ART-based hybrid drugs were developed and their activity profile was characterized against drug-sensitive and drug-resistant *Plasmodium falciparum* parasites. Two hybrids were found to display parasite growth reduction, stage-specificity, speed of activity, additivity of activity in drug combinations, and stability in hepatic microsomes of similar levels to those displayed by dihydroartemisinin (DHA). Conversely, the rate of chemical homolysis of the peroxide bonds is slower in the hybrids than in DHA. From a mechanistic perspective, heme plays a central role in the chemical homolysis of peroxide and in inhibiting heme detoxification and disrupting parasite heme redox homeostasis. The hybrid exhibiting slow homolysis of peroxide bonds was more potent in reducing the viability of ART-resistant parasites in a ring-stage survival assay than the hybrid exhibiting fast homolysis. However, both hybrids showed some limited activity against ART-induced quiescent parasites in the quiescent-stage survival assay. Our findings are consistent with previous results showing that slow homolysis of peroxide-containing drugs may retain activity against proliferating ART-resistant parasites. However, our data suggest that this property does not overcome the limited activity of peroxides in killing non-proliferating parasites in a quiescent state.

**GRAPHICAL ABSTRACT:** 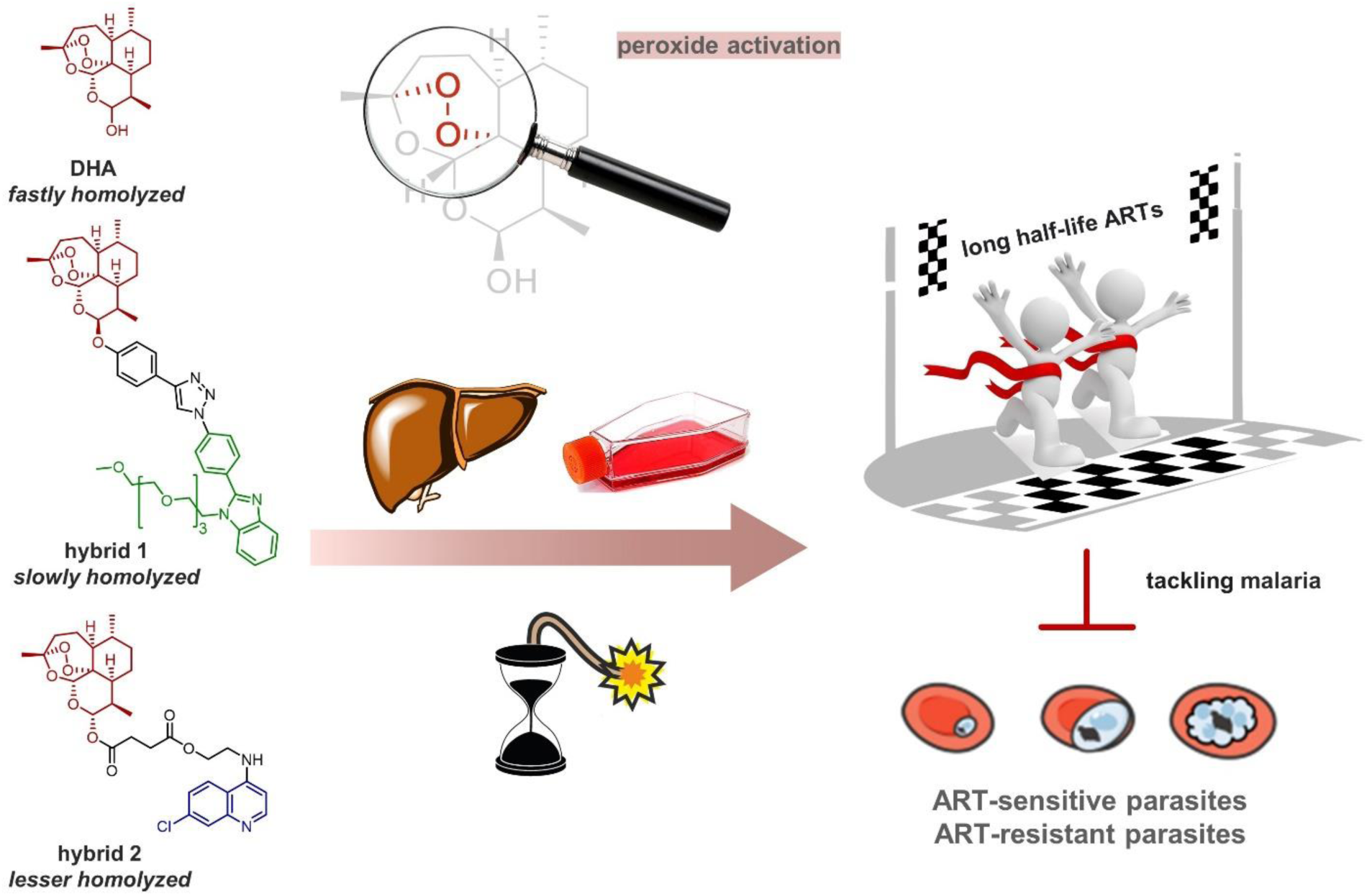

Hepatic and cell-host-mediated metabolism are responsible for short plasma half-lives of antimalarial artemisinins (ARTs), illustrated here by dihydroartemisinin (DHA). ART-based hybrid drugs that overcome rapid degradation can facilitate activity against ART-resistant parasites, as illustrated by hybrid **1**.

## 1. INTRODUCTION

Artemisinins (ARTs) are the core components of antimalarial therapy at the present time [1, 2]. However, their role as antimalarial drugs has been threatened by the rise of parasites with a decreased susceptibility to ART treatment [3]. The main phenotypic hallmark of ART-resistant parasites is a prolongation of their ring stages during asexual blood stage development [4] and the ability of part of these parasites to enter quiescence upon exposure to ARTs [5]. This process leads to delayed-clearance phenotype (DCP) parasites, whose main genotypic hallmark is a polymorphism in the *kelch-13* gene [6–9]. The implementation of long-lasting drugs as partners in ART-based combination therapy (ACT) has helped mitigate the spread of DCP parasites. Of concern is the observation that ART-resistant parasites in a quiescent state display low cellular metabolism [9], which is phenotypically characterized by reduced susceptibility to heme detoxification suppressors like amodiaquine (AQ) and piperaquine, which are often deployed in ACT [10, 11]. Only a limited number of drugs are active against quiescent parasites, including mefloquine (MFQ) and atovaquone [11].

The peroxide bonds of clinically employed ART-derived drugs can be homolyzed in plasma, blood, and tissues. While metabolism to inactive molecules occurring in plasma and liver can contribute to drug degradation [12–15], heme is by far the most compelling agent of peroxide activation. As a reducing agent of peroxide, Fe(II)-heme plays a key role by leading to the formation of a *C*-centered radical able to alkylate heme and proteins [16]. It is this activation pathway that has been shown to be responsible for the antiplasmodial activity of peroxide-based antimalarials [16–19]. Despite some subtle differences in the peroxide homolysis products among the ART-derived drugs used clinically [15], the elimination half-life (t_1/2_) of all these drugs in the plasma is short and considered to be the bottleneck [12, 20].

To overcome this drawback, peroxide drugs with extended t_1/2_ have been developed by replacing the 1,2,4-trioxane present in the ART with a 1,2,4-trioxolane (ozonides) or a 1,2,4,5-tetraoxane. Studies have shown that ozonide OZ439 (t_1/2_ >42 h) and tetraoxane E209 (t_1/2_ >30 h) have a longer half-life than ozonide OZ277 (t_1/2_ ∼2-4 h) and DHA (t_1/2_ ∼1-3 h) [21–23]. While all these peroxide-based antimalarials are known to kill parasites by the same mechanism of radical formation [16, 24], extending the t_1/2_ of a peroxide drug can modify the parasite’s susceptibility to treatment. For instance, OZ439, which has a much higher t_1/2_, is capable of killing ART-resistant parasites more effectively than OZ277, while DHA fails to kill them [22]. The underlying reason for this efficacy is attributed to the slower metabolic degradation of OZ439, which enables this drug to surpass the parasites’ capability to withstand the drug-induced oxidative stress and to alkylate protein unaffected by ARTs [24].

For drugs derived from 1,2,4-trioxolane or 1,2,4,5-tetraoxane, half-lives can be increased by enhancing the hydrophobicity of the molecule in order to improve the drug-like properties, such as logD, solubility, and metabolic stability, thereby optimizing the pharmacokinetic profile [25, 26]. Another approach to overcome the shortcomings of ARTs is the development of ART-based hybrid compounds containing a second pharmacophore. Examples of such ART-based hybrids include ones in which the second pharmacophoric group consists of 4-aminoquinolines, facilitating drug penetration inside the parasite and inhibiting heme polymerization synergistically with heme alkylation [27–29]. ARTs have also been conjugated with vinyl phosphonates to inhibit cysteine proteases and with peptidomimetics to inhibit proteasomes [29–31]. These hybrids display improved activity against drug-resistant parasites, even when the second moiety has no obvious activity on its own, ultimately indicating that a second pharmacophoric group and a chemical linker might induce conformational changes around the peroxide bond, protecting it from a rapid reaction [32–34]. This evidence supports the hypothesis that the chemical design of ART-derived drugs can extend their half-lives beyond those of the parental ARTs.

The rate of homolysis of peroxide bonds assayed in red blood cells, also referred to as cell-host-mediated degradation [13, 33, 35], arguably explains the effectiveness of ozonides against ART-resistant parasites [23, 36]. This kind of knowledge is crucial for understanding the susceptibility profile of clinically employed ART-derived drugs [13,33,35]. However, little direct, quantitative evidence of cell-host-mediated degradation in ART-based hybrid drugs has been reported. Here, we characterize the susceptibility of sensitive and resistant parasites to treatment with ART-based hybrid drugs. The results reveal the activity of the hybrid drugs to be superior to that of the parental DHA against ART-resistant *P. falciparum* parasites. Additionally, we assess the hybrid drugs’ cell-host-mediated degradation and chemical stability in hepatic microsomes. The hybrid exhibiting the greatest stability against cell-host-mediated degradation was also the most potent antimalarial agent *in vitro*, and the most stable hybrid in the microsomes was found to be the most efficacious treatment in mice. In addition, we examined the steps underlying the reductive bioactivation of peroxide by heme and their effect in killing quiescent ART-resistant parasites.

## 2. MATERIALS AND METHODS

### 2.1 Drugs

Atovaquone (ATO), amodiaquine (AQ), chloroquine (CQ), dihydroartemisinin (DHA), and mefloquine (MFQ) were obtained from Sigma-Aldrich (St. Louis, MO, USA). ART-based hybrid drugs LH70 (“hybrid **1**”) and 163A (“hybrid **2**”) were synthesized as described elsewhere [29, 34]. SYBR Green I nucleic acid gel stain was purchased from Thermo Fischer Scientific (Waltham, USA). All other general chemicals and solvents were of analytical or HPLC grade.

### 2.2 Culture of *P. falciparum*

ART-susceptible strains NF54, D10, 3D7, and W2 of *P. falciparum* were grown using standard methods. These parasite strains were maintained at 5% hematocrit (human type O- or A-positive red blood cells [RBC]) in RPMI-1640 Medium (EuroClone, Milan, Italy) supplemented with 1% AlbuMax (Invitrogen, Milan, Italy), 2 mM L-glutamine (Euroclone), 20 mM HEPES (Euroclone), and 0.37 mM hypoxanthine (Sigma-Aldrich, St Louis, USA). For parasite growth and experiments, the cultures were maintained at 37°C in a standard gas mixture composed of 1% O_2_, 5% CO_2_, and 94% N_2_. For the ring-stage survival assay (RSA) and quiescent-stage survival assay (QSA), ART-sensitive (F32-TEM) and ART-resistant (F32-ART) strains were used. F32-ART is a laboratory strain resistant to ARTs obtained after several years of sequential and increasing artemisinin pressure, while F32-TEM is its susceptible isogenic twin [5]. F32-ART carries a M476I mutation on the *pfk13* gene that is responsible for its ART resistance [6]. These two strains were cultured in RPMI-1640 Medium (Dutscher, Bernolsheim, France) at 2% hematocrit in human RBC (EFS, French blood bank, Toulouse, France) and 5% human serum (EFS, French blood bank, Toulouse, France) at 37°C in a 5% CO_2_ humidified atmosphere.

### 2.3. Antiplasmodial activity against ART-susceptible *P. falciparum* parasites

Each compound was dissolved in dimethyl sulfoxide (DMSO) and diluted in RPMI-1640 Medium into seven different concentrations, which were tested against CQ-susceptible strains (NF54 or D10) or a CQ-resistant strain (W2). Each well of 96-well plates was filled with 100 µL drug and 100 µL parasitized RBC (asynchronous parasite culture), which were mixed to obtain a final parasitemia of 1.0% to 1.5% and 1.0% hematocrit. Plates were incubated for 72 h at 37°C in a standard gas mixture, then parasite growth was determined by measuring the activity of the parasite lactate dehydrogenase (pLDH) [37]. Briefly, at the end of incubation, the cultures were carefully resuspended, and 20 μL aliquots were removed and added to 100 μL Malstat reagent in a 96-well microplate. The Malstat reagent is made of 0.125% TritonX-100, 130 mM L-lactate, 30 mM Tris buffer, and 0.62 μM 3-acetylpyridine adenine dinucleotide. After that, 20 μL 1.9 μM nitro blue tetrazolium (NTB) and 0.24 μM phenazine ethyl sulphate were added to the plate. NBT was reduced to blueformazan and spectrophotometrically read at OD 650 nm using a Synergy 4 microplate reader (BioTek, Santa Clara, USA). The concentration at which the drugs were able to inhibit 50% parasite growth (IC_50_) was calculated using the inhibitory effect sigmoid Emax model, estimating the IC_50_ value through nonlinear regression using a standard function of the software package R (ICEstimator version 1.2). IC_50_ values were expressed as means of two to three independent experiments, with each drug concentration in duplicate.

### 2.4 Antiplasmodial activity of drug combinations

The activity of the drug combinations against the *P. falciparum* NF54 strain was determined as described above. In each experiment, drugs were tested alone and in combination. Twofold serial dilutions of the drug solutions were made using fixed molar ratios of 1:1, 1:3, and 3:1 for each combination and using two technical replicates (ranging from 200 to 1.0 nM). Parasite growth was determined by the pLDH method, as described above. IC_50_ values for each drug alone or in combination were determined and an isobologram analysis was performed by plotting the fractional inhibitory concentrations (FIC) for each combination and its component drugs separately. Individual FIC values were calculated as the IC_50_ of the drug combination divided by the IC_50_ of the drug in isolation. A dashed line was plotted to indicate any additive effect and to distinguish antagonism (above the dashed line) from synergism (below the dashed line). Three independent experiments were performed, each one containing two replicates. Each experiment was used to represent an isobologram analysis (mean and 95% confidence interval [CI]).

### 2.5 Ring-stage-specific drug activity

The NF54 strain of *P. falciparum* was employed to test the drugs’ activity against ring stages and asynchronous parasite culture (all stages). The ring stages were obtained by synchronization with D-Sorbitol using a standard method [38]. Each well of 96-well plates was filled with 100 µL each drug and 100 µL culture containing ring-stage or asynchronous parasites, which were mixed to obtain a final parasitemia of 1.0% to 1.5% and 1.0% hematocrit. Plates were incubated for 72 h at 37°C in a standard gas mixture. Afterwards, plates were centrifuged, cell pellets were carefully washed twice with complete medium without drugs, and the plates were returned to the incubator for an additional 66 h. Plates were processed using the pLDH method and IC_50_ values were calculated as described above. At least two to three independent experiments were performed, using two technical replicates.

### 2.6 Exposure time dependence of drug activity

Each well of 96-well plates was filled with 100 µL drug and 100 µL culture containing ring-stage *P. falciparum* (NF54 strain) parasites, which were mixed to obtain a final parasitemia of 1.0% to 1.5% and 1.0% hematocrit. Plates were incubated at 37°C in a standard gas mixture. After 3 h and 6 h, plates were centrifuged, cell pellets were washed as described above then returned to the incubator for an additional 69 h and 66 h, respectively. In parallel, the same experiment was left without the washing step, enabling continuous drug exposure throughout. Plates were processed using the pLDH method and IC_50_ values were calculated as described above. Three independent experiments were performed using two technical replicates.

### 2.7 Speed of drug activity

Each well of 96-well plates was filled with 100 µL drug and 100 µL culture containing asynchronous *P. falciparum* (NF54 strain) parasites, which were mixed to obtain a final parasitemia of 1.0% to 1.5% and 1.0% hematocrit. The plates were incubated for 24, 48, and 72 h at 37°C with a standard gas mixture. DHA (a fast-acting drug) was used as a positive control and ATO (a relatively slow-acting drug) was used as a negative control [39]. Parasite viability was measured using the pLDH method and IC_50_ values were calculated as described above. Three independent experiments were performed using two technical replicates.

### 2.8 Cell-host-mediated drug degradation

Each well of 96-well plates was filled with 125 µL drug (1000 nM) and 125 µL uninfected RBC (uRBC), which were mixed to obtain a final drug concentration of 500 nM and 2.5% or 1.0% hematocrit. The plates were incubated at 37°C in the standard gas mixture. After each timepoint (0.16, 6, and 24 h), supernatants were carefully harvested and stored at −80°C until use. Untreated uRBC incubated at the same times were used as negative controls and DHA and AQ were employed as standard drugs. Next, supernatants were diluted in complete medium in six different dilutions and were aliquoted in new plates. The antimalarial activity of these supernatants and of freshly dissolved drugs was determined in an asynchronous parasite culture of *P. falciparum* (3D7, D10, and W2 strains) for 72 h using the same method described above, and parasite viability was assessed by the pLDH and SYBR Green I methods. Only the experiments where the IC_50_ values of the supernatants harvested at 0.16 h were the same as those of the respective fresh drugs were considered for determining equivalent drug concentration. Assuming the supernatant harvested at 0.16 h had a drug concentration of 500 nM, an equivalent drug concentration in the supernatants was calculated as 500/IC_50_ value for each timepoint. Decay in drug concentration was estimated by using the normalized slope values calculated from linear regression fits using the slope values of a standard curve as reference. Three independent experiments were performed using two replicates.

### 2.9 *In vitro* metabolic stability assay in microsomes

Metabolic stability studies were conducted in human, mouse, and rat liver microsomes using a single-point (30 min.) study design [40]. Briefly, 1.0 μM of the compound was incubated with 0.4 mg/mL microsomes in 0.1 M phosphate buffer (pH 7.4). The reactions were quenched by the addition of ice-cold acetonitrile containing an internal standard (carbamazepine, 0.0236 μg/mL). The samples were centrifuged, and the supernatant was analyzed by LC−MS/MS using an Agilent Rapid Resolution HPLC with an AB SCIEX 4500 MS instrument (Santa Clara, USA). The percentage of compound that remained was calculated by using the internal standard corrected peak areas at 30 min, compared to those at t_0_. Propranolol, midazolam, and MMV390048 were used as assay controls (data not shown).

### 2.10 Chemosensitivity on ART-resistant parasites using standard assay

Compounds were dissolved in DMSO and diluted in RPMI-1640 Medium into five different concentrations (0.1–1000 nM). Each well of 96-well plates was filled with 100 µL drug at each concentration and 100 µL synchronized ring-stages of the ART-resistant strain F32-ART or its ART-susceptible twin strain F32-TEM at final parasitemia of 1% and 2% hematocrit. Plates were incubated for 48 h at 37°C and 5% CO_2_. The drugs were the washed out in 1X PBS (Sigma-Aldrich), the plates were frozen and thawed, and then the lysed parasites were transferred to black 96-well plates. SYBR Green I (Fisher Scientific, Illkirch, France) at a 2X concentration in a lysis buffer (20 mM TRIS base, pH 7.5, 20 mM EDTA, 0.008% w/v saponin, 0.08% w/v Triton X-100) was then added and the plates were incubated for 1 h at room temperature in the dark. Fluorescence was measured at 485 nm excitation and 528 nm emission in a VICTOR® Nivo™ plate reader (Perkin Elmer, Waltham, USA) and the IC_50s_ values were then calculated using Prism 7 (GraphPad, San Diego, USA).

### 2.11 *In vitro* recrudescence assay

F32-ART and F32-TEM parasites were synchronized at the ring stage by D-sorbitol treatment, adjusted to 3% parasitemia and 2% hematocrit, and treated with 700 nM of each drug for 48 h in a 6-well plate, as 700 nM concentration corresponds to the plasma peak of DHA in patients [41]. Parasites were washed with RPMI-1640 Medium before being placed in drug-free culture conditions with 10% human serum. Parasitemia was determined daily by Giemsa-staining of blood smears until the cultures reached their initial parasitemia, defined as the recrudescence day. If no parasite recrudescence was observed for up to 30 days, the experiment was stopped. Two independent experiments were performed.

### 2.12 Ring-stage survival assay (RSA^0-3h^)

Early ring-stage parasites (0–3 h post-invasion) of F32-ART and F32-TEM strains highly synchronized by Percoll-Sorbitol treatment were exposed to 700 nM of the drugs or 0.1% DMSO (control) for 6 h. Each drug was tested in duplicate. Parasite pellets were then washed in RPMI-1640 Medium before being placed in drug-free culture conditions with 10% human serum and incubated for 66 h. Survival rates were assessed microscopically by two independent microscopists by counting the parasites in 10,000 RBCs. Three independent experiments were performed.

### 2.13 Quiescent-stage survival assay (QSA)

The QSA was performed as previously described to determine the drug activity on quiescent ART-resistant parasites [11]. F32-ART culture parasites were synchronized at the ring-stage and adjusted to 3% parasitemia and 2% hematocrit. The parasites were then exposed for 6 h to 700 nM DHA in order to mimic the plasma peak of DHA in patients [41] and to induce quiescence (Figure 4C, conditions A and B) or a mock condition (Figure 4C, condition C). After the washing steps, quiescence was maintained for 48 h in parasites treated with either DHA alone (Figure 4C, condition A) or DHA plus 700 nM of the drug to be tested (Figure 4C, condition B). At the end of treatment, parasites from each of the three conditions were washed with RPMI-1640 Medium and replaced in new wells in drug-free culture medium with 10% human serum. Parasitemia was determined daily by thin blood smears until the cultures reached their initial parasitemia (3%), which was defined as the recrudescence day [11]. The recrudescence capacity of parasites exposed in conditions A and B was compared to determine whether the drug was active on quiescent parasites. Condition C was designed to check that the concentration at which the drug was tested acted on proliferating parasites. If no parasite recrudescence was observed, the experiment was stopped.

### 2.14 Fractionation assay of parasite-heme-derived species

Compounds were dissolved in DMSO and diluted in RPMI-1640 Medium into four different concentrations (1.56, 6.25, 25, and 100 nM). Each well of 24-well plates was filled with 1.0 mL ring stages of 3D7 strain *P. falciparum* at final parasitemia 5% and hematocrit 4%. Plates were incubated for 32 h at 37°C in standard gas mixture. Cell pellets were then washed, the supernatant was discarded, and the pellets were resuspended in a lysing solution (1.0% w/v saponin). The resulting suspension was centrifuged, the supernatant was discarded, and the pellets were washed twice with phosphate buffered solution (PBS). The resulting pellets were then stored at −80°C until analysis. Parasite-derived heme species (hemoglobin, free heme, and hemozoin) were determined by successive fractionation assays in a 96-well plate format using a standard method [42]. The relative amount of each species was determined by summing all three species (hemoglobin, free heme, and hemozoin). Three independent experiments were performed using two replicates of each drug concentration.

### 2.15 β-hematin inhibitory activity (BHIA)

The compounds were dissolved in DMSO and diluted in DMSO to six different concentrations (1.0–32 mM). Chloro-hemin (Fe^III^[PPIX]Cl) (BioXtra, Sigma-Aldrich) was first dissolved in 0.1 M NaOH then suspended in an 80:20 v/v mixture of DMSO and propionate buffer (0.1 M, pH 5.6) to a stock solution at 8 mM. Reduced glutathione (GSH) was dissolved in 0.1 M NaOH and then adjusted to 10 mM using propionate buffer (0.1 M, pH 5.6). For BHIA under the reducing condition (R-BHIA), hemin (10 µL) and GSH (10 µL) were distributed in a 96-well plate, homogenized, and the plates were sealed using a thin plastic film and incubated for 1 h at 37°C. Then, 20 µL drug was added and the plates were incubated for 2 h at 37°C. The plastic film was removed after drug incubation, 160 µL propionate buffer (1.0 M, pH 5.6) was added, and plates were incubated for 18 h at 37°C. BHIA under the oxidizing condition (O-BHIA) was prepared in parallel without adding GSH. Controls included wells without drugs and wells containing PBS 1x instead of propionate buffer. Afterwards, the plates were centrifuged, the supernatant was discarded, and the pellets of β-hematin crystals were washed twice with pure DMSO and then water. The resulting β-hematin crystals were then dissolved in 200 µL 0.1 M NaOH and diluted to a 1:8 ratio. Aliquots were then transferred into a new plate and read by UV-visible absorbance at 405 and 450 nm. Percent of BHIA was calculated in comparison to untreated wells (no drugs) and IC_50_ was then calculated. Two independent experiments were performed using three replicates of each drug concentration [43].

### 2.16 Statistical analysis

Data were typically presented as median ± standard error of the mean (SEM) or mean ± standard deviation (SD). The drug concentration required to inhibit 50% parasite growth (IC_50_) was calculated by nonlinear curve fitting of log-transformed and normalized data using Prism 8.4.0 or ICEstimator 1.2 (http://www.antimalarial-icestimator.net/index.htm). Statistical analyses were performed using Prism 8.4.0. Any outlier was identified using Grubbs’ test and excluded from the data analysis. Statistical significance was assessed by nonparametric and unpaired Student’s *t* test for sample comparison or two-way ANOVA for multiple comparisons (corrected by Bonferroni post-test) as indicated in each figure. RSA values were determined by the Mann-Whitney unpaired *t*-test. Statistical significance was set at *p* <0.05. Results from the recrudescence assay and QSA were interpreted using Kaplan-Meier analysis, considering censored data, and statistical significance was assessed by the Log-rank Mantel-Cox test using Prism 7.

## 3. RESULTS

### 3.1 Chemical design

We previously set up a medicinal chemistry program to synthesize libraries of ART-based hybrid drugs with different pharmacophoric structures covalently bound to ART through a variety of chemical linkers [29,33,34,44,45]. The study of pharmacophores, linkers, and substituents revealed that benzimidazoles with a non-cleavable triazole linker afforded ART-based hybrids a potent and efficacious antimalarial property as well as stability and auto-fluorescence, suitable for tracking the location of the drugs in living cells. ART-benzimidazole hybrid **1** was the most potent among them and it was chosen as a representative non-cleavable hybrid drug [34]. We also deployed ART-quinoline hybrid **2** as a representative cleavable hybrid drug. Hybrid **2** combines an ART moiety with a quinoline group via a cleavable ester linker, which has shown potent activity against *P. falciparum in vitro* and efficacy in *P. berghei*-infected mice of a higher level than ART but lower than hybrid **1**. Moreover, the chemoproteomics of the compound (**2**) have revealed a capacity to alkylate proteins comparable to ARTs [29,32]. Importantly, these two hybrids have displayed superior antimalarial activity against CQ-resistant parasites than their two separated moieties when administered as a drug combination (ART plus quinoline/benzimidazole). Their chemical structures and key chemical and pharmacological features are shown in **Figure 1**.

**Figure 1.**
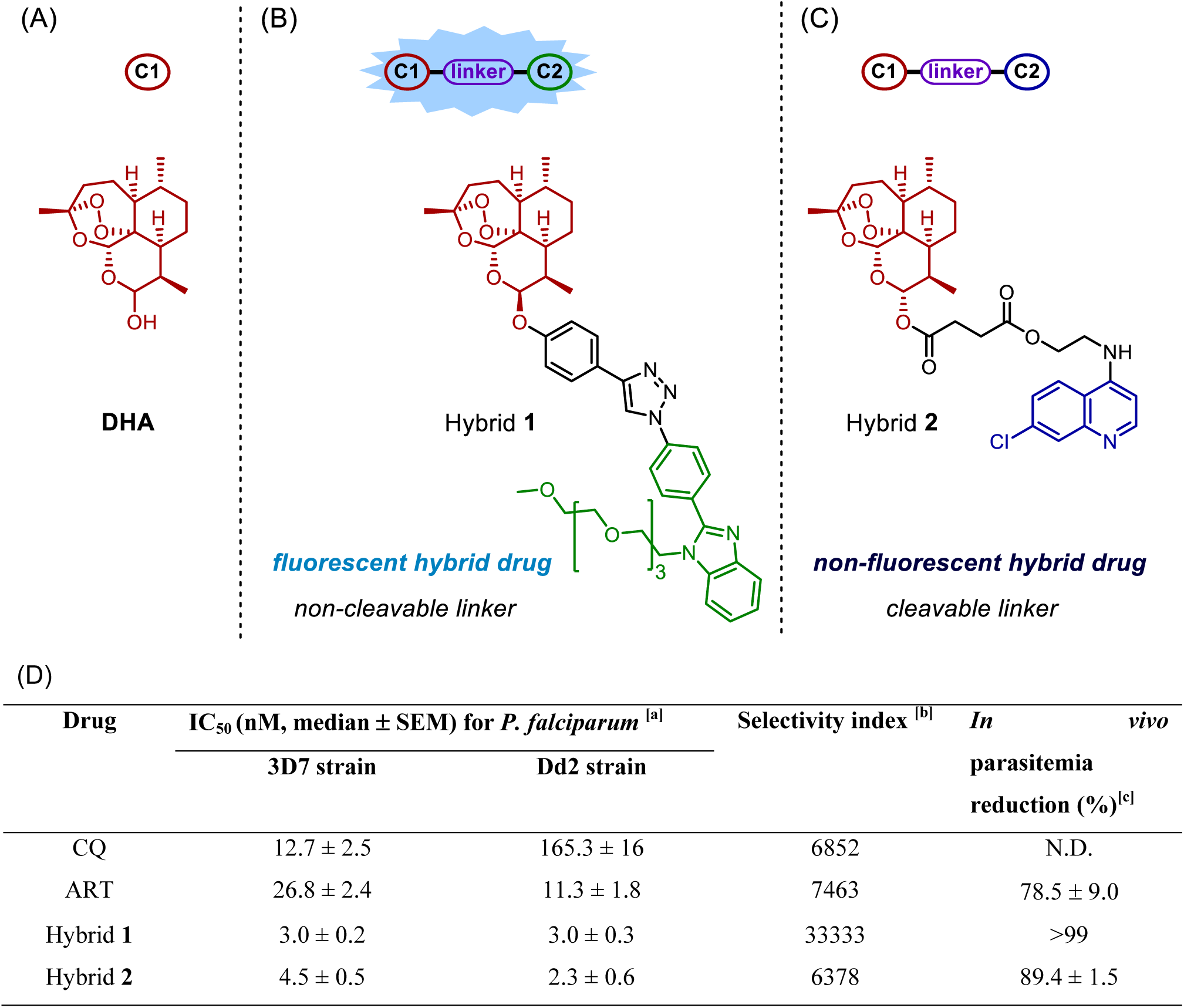
Chemical structures of the ART-based hybrid drugs studied here and their antimalarial activity. Panel A shows the structure of peroxide trioxane dihydroartemisinin, and the pharmacophore groups employed for the design of ART-based hybrid drug **1** (panel B) and **2** (panel C). Panel D shows a table summarizing the antimalarial activity of each drug (values taken from [29, 32, 34]). Footnotes for Table: ^[a]^ Assay determined in asynchronous culture and activity measured by SYBR Green I; ^[a]^ Indices were determined by CC_50_ / IC_50_, where CC_50_ was assayed in a J774 cell line. ^[a]^ Efficacy determined in *P. berghei*-infected mice using Peters test. SEM = standard error of the mean; CQ = chloroquine; ART = artemisinin; N.D. = not determined.

### 3.2 Hybrids can efficiently kill young parasite stages with exposure time dependency

We studied the stage-specific susceptibility of drug activity against the asexual blood stages of *P. falciparum* parasites using the NF54 strain (**Figure 2A**). To determine the specificity of the drugs against ring stages, we incubated parasites with the compounds for 6 h, washed out the drugs, and measured drug activity on parasite growth 66 h later [6,47]. Of note, we assured the complete removal of any residual and unbound drugs in the cell supernatant after the washing out steps and replacement of culture medium by quantification of the autofluorescence of hybrid **1** in the cell supernatant (Figure S1).

**Figure 2.**
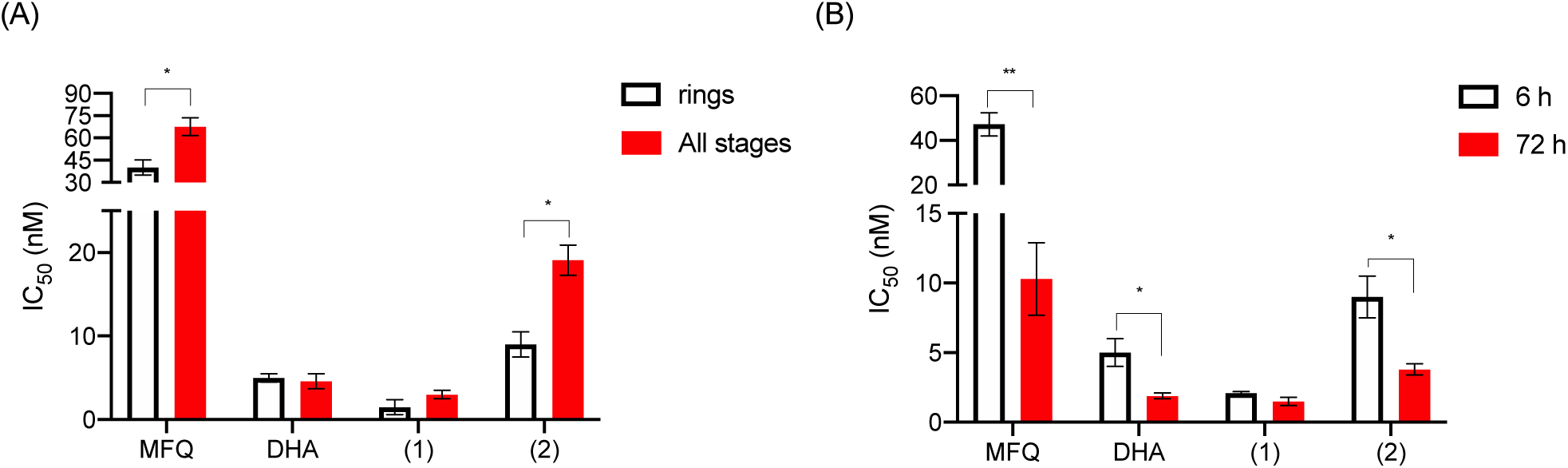
Hybrids kill young parasite stages with exposure time dependency. Panel A shows the ring stage-specific susceptibility and panel B shows the exposure time dependency of the drug activity against NF54 strain of *P. falciparum.* In panel A, parasites at all stages (ring stages or asynchronous) were incubated in the presence of drugs for 72 h. In panel B, ring-stage parasites were incubated in the presence of drugs for short-pulse (6 h) or standard (72 h) incubation. In both cases parasitemia was assessed at 72 h. Parasite viability was measured by pLDH, and IC_50_ values were calculated. Values are shown as the median and SEM (error bars) of three independent experiments, using each concentration of compounds in duplicate. \**p* <0.05; \*\**p* <0.01 were significantly different by Student’s *t* tests. (1) = hybrid **1**; (2) = hybrid **2;** MFQ = mefloquine; DHA = dihydroartemisinin; SEM = standard error of the mean. **Figure S2** and **Table S1** (supporting material) show the detailed results.

DHA was equally potent in killing ring-stage and asynchronous *P. falciparum* parasites. In contrast, MFQ was more potent at killing ring-stage than asynchronous parasites. Hybrid **1** demonstrated similar effectiveness at killing young parasite stages and asynchronous parasites **(Figure 2A)**, but less activity against asynchronous parasites. This was attributed to the presence of the 4-aminoquinoline component, which is also present in CQ [47] and is known to be less effective at killing late trophozoites and schizonts. Overall, our results of stage-specific susceptibility for peroxide-based drugs (DHA, hybrids) and MFQ are broadly consistent with the literature [47].

Given that young parasite stages were susceptible to both hybrids, we sought to understand the exposure time dependency of the drug activity. Ring-stage parasites were exposed to three different conditions of drug treatment: short-time exposure (3 h and 6 h) and standard-time exposure (72 h). In all the experiments, parasite growth reduction was assessed after 72 h of incubation. Parasites showed lower sensitivity against MFQ, DHA, and hybrid **2** after 3 h and 6 h exposure than after 72 h exposure. For hybrid **1**, no statistical difference was found according to duration of treatment **(Figure 2B)**. Despite some dissimilarities in the exposure time to effectively kill parasites, most peroxide-containing drugs tend to be associated with a rapid onset of parasite viability reduction, namely, the parasite viability is already reduced when assessed after 24 h incubation with drugs. The IC_50_ values determined after 24 h of treatment were of similar range as the values after 48 h and 72 h (Figure S3). These data confirm that both hybrids can kill parasites with a speed of action comparable to that of parental DHA.

### 3.3 Hybrids are useful in drug combinations

We assessed hybrids in drug combination studies. To this end, CQ (a heme detoxification suppressor) and MFQ (a pleiotropic agent), which are two reference antiplasmodial drugs, especially in regions with a low prevalence of ART-resistant parasites, were chosen for the combination studies. Assays were performed using different molar ratios of the drugs and against ring stages of the NF54 strain of *P. falciparum* cultivated for 72 h (Figure S4, Table S2). We observed nondetrimental interactions towards additivity between hybrid **1** and CQ, hybrid **1** and MFQ, and hybrid **2** and MFQ, while hybrid **2** showed an antagonistic effect against CQ. The detrimental interaction of hybrid **2** and CQ is an exception worthy of further interpretation. Both are 4-aminoquinoline-based drugs, which could lead to competition for the mechanism to inhibit parasite growth. Overall, most drug combinations demonstrated an additive effect with respect to their ability to inhibit parasite growth. This is in close agreement with the literature for ACT, in which additive effects have been observed [43], while a high level of synergy has only been found for atovaquone plus proguanil [48].

### 3.4 Hybrids have slower cell-host-mediated degradation than DHA

We examined whether the hybrids underwent any cell-host-mediated degradation, as is observed for ARTs [13,35,36]. We assayed this by incubating the drugs at 500 nM in uninfected red blood cells (uRBC) [33]. The supernatants were harvested at different time points (10 min., 6 h, 24 h) and antiplasmodial activity was determined for them in comparison with freshly added drugs. We detected residual amounts of lysed uRBC at high drug concentrations (700 and 1000 nM) and at an incubation time of 48 h, which could potentially interfere with an accurate measurement of antiplasmodial activity. Given the short-lived property of ARTs, harvesting supernatants after drug incubation up to 24 h was adequate for estimating cell-host-mediated degradation. AQ was used as a long-lasting drug [47], while DHA was used as a short-lived drug [13] (**Figure 3A**).

**Figure 3:**
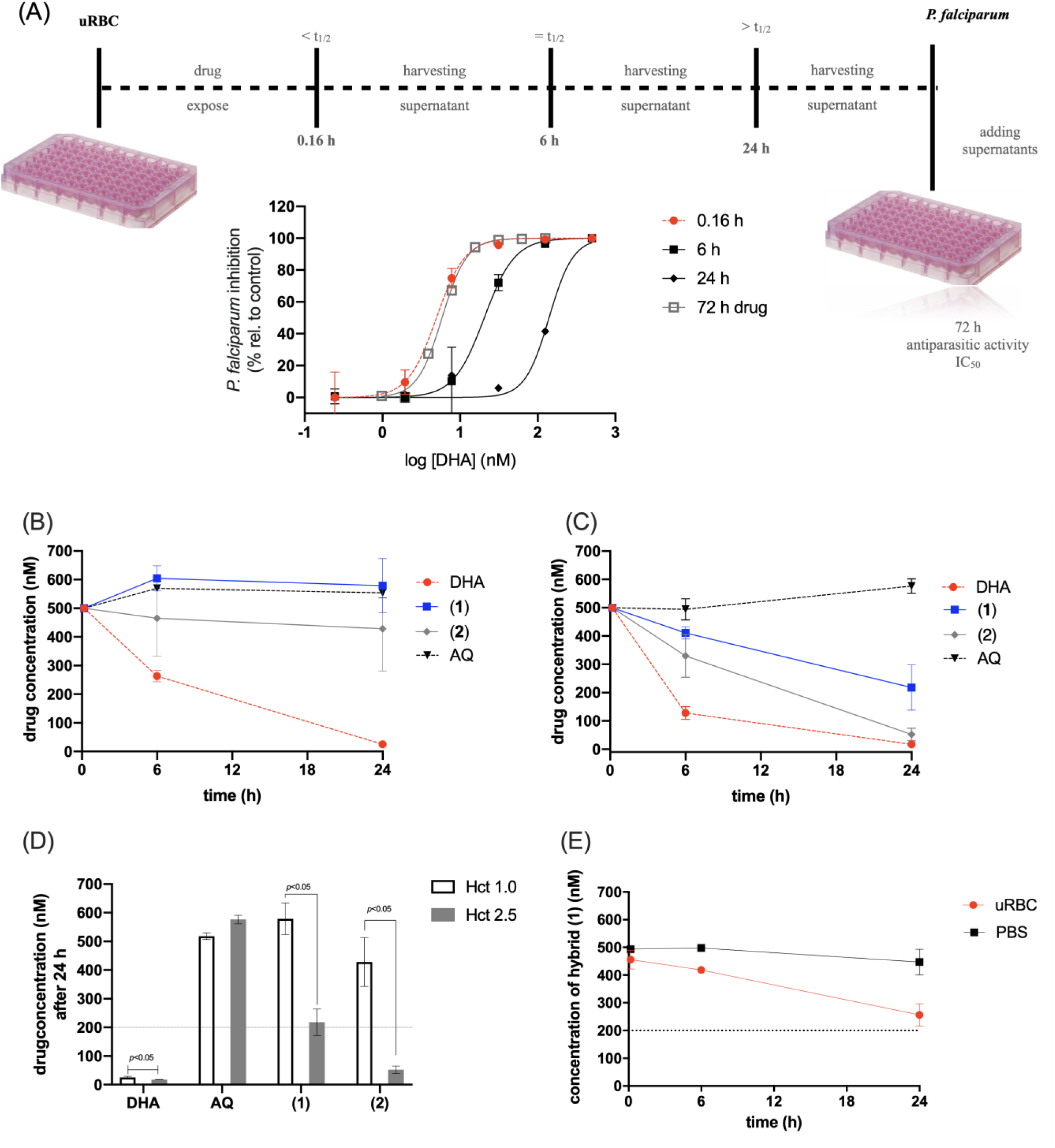
Hybrids have slower cell-host-mediated degradation than DHA. Panel (A) shows the experimental design to assay cell-host-mediated drug degradation and the analysis to calculate the chemical stability of the drugs. Panel (B) shows cell-host-mediated degradation at low hematocrit (1.0%). Panel (C) shows degradation at high hematocrit (2.5%). Panel (D) shows the drug concentration of supernatants harvested at 24 h. Panel (E) shows the concentration of hybrid **1** in the supernatants of uRBC and PBS determined by HPLC. uRBC were exposed to 500 nM compound and the supernatants were harvested at the indicated time (0.16, 6, and 24 h). Antiplasmodial activity of the supernatant and of freshly diluted drugs were assayed against the asynchronous 3D7 strain of *P. falciparum*. The method to estimate the (equivalent) drug concentration is described in the experimental section. Indicated values were significantly different by Student *t* tests. AQ = amodiaquine. DHA = dihydroartemisinin. PBS = phosphate-buffered saline. **Figure S5** (supporting material) shows the detailed results.

**Figure 4:**
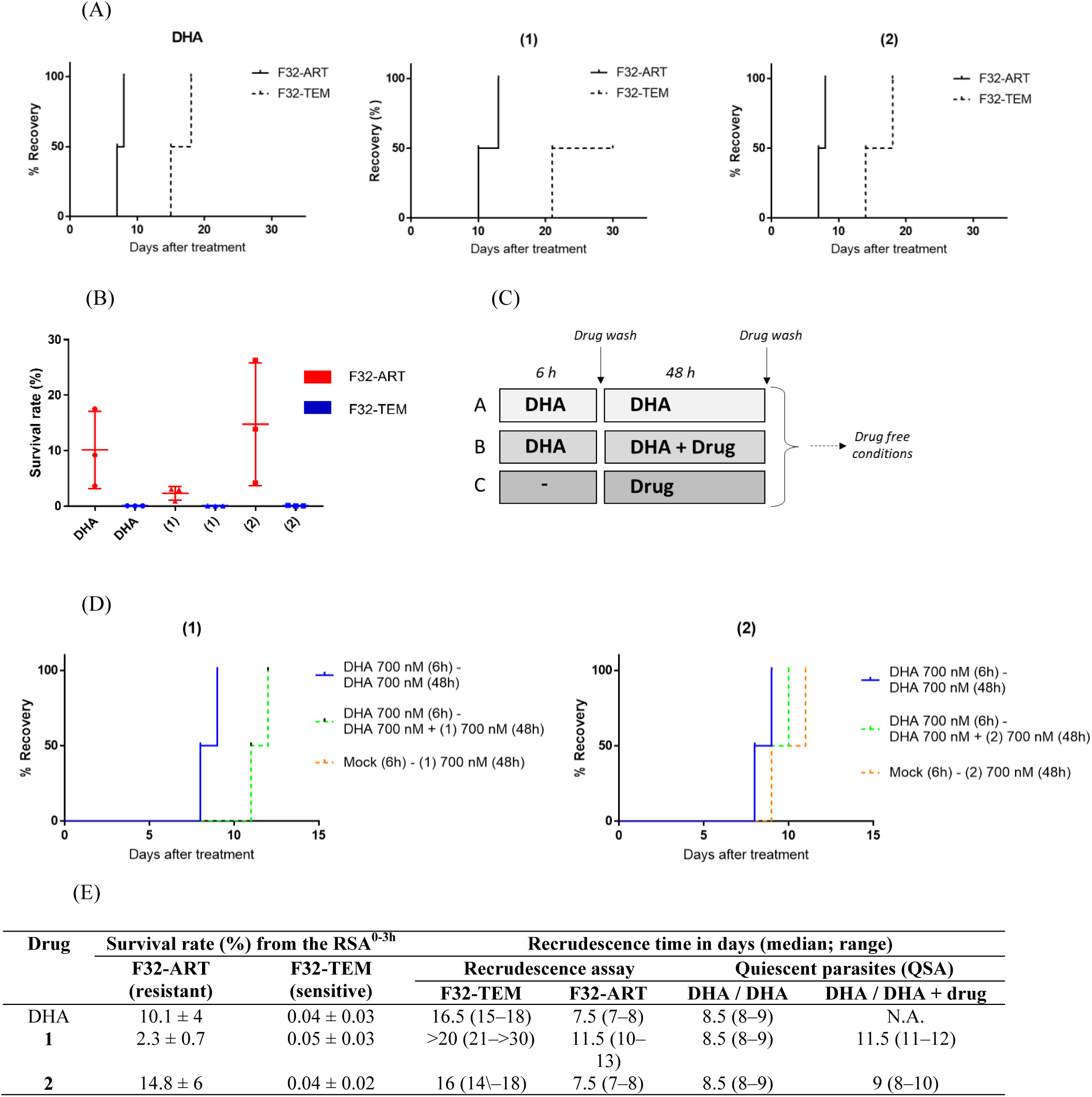
Hybrids exhibit enhanced killing properties against ART-resistant *P. falciparum* parasites. Panel A shows that K13 mutation conferred cross-resistance to DHA and hybrid **2** but less so to hybrid **1** treatment, as defined in the recrudescence assay with ring-stage parasites cultured under treatment for 48 h. The final event was defined as the time necessary for the treated cultures to reach their initial parasitemia, by monitoring recrudescence daily. Data were censored if no recrudescence was observed at day 30. Two independent experiments were performed in parallel for F32-ART and F32-TEM lines in the same conditions to generate paired results. Panel B shows the percent survival values from drug-treated mutant-K13 (F32-ART) and wild-type-K13 (F32-TEM) lines in the RSA^0-3h^. Drugs were incubated for 6 h, compounds were washed out, and parasite cultures were allowed to grow for 66 h. Viable parasites were quantified by Giemsa-stained smears read by two microscopists and their numbers normalized to values for the DMSO control. Data correspond to three independent experiments. Statistical significance was determined by Mann-Whitney unpaired t-test. Panel C shows the experimental design for the quiescent-stage survival assay (QSA). Panel D shows that quiescent parasites are more susceptible to treatment with hybrid **1** than to treatment with DHA or hybrid **2**, as defined by Kaplan-Meier analysis of recrudescence in the QSA. Data correspond to two independent experiments. For hybrid (1), the orange curve is superposed to the green one. Panel E shows a table summarizing all the values. In all experiments, drugs were added at a concentration of 700 nM. N.A. = not applicable; DHA = dihydroartemisinin; RSA^0-3h^ = ring-stage survival assay; QSA = quiescent-stage survival assay.

At a low hematocrit level (1%), DHA-derived supernatants presented a subtle (twofold) loss in their antiplasmodial activity when harvested at 6 h and a significant loss (as high as 20-fold) at 24 h **(Figure 3B**). In contrast, AQ-derived supernatants for different harvesting times displayed equipotent antiplasmodial activity. Under these conditions, neither hybrid showed any measurable drug degradation, even at 24 h. To subsequently confirm that the loss in supernatant activity was due to cell-host-mediated degradation, which is intrinsically dependent on the levels of bioavailable heme, we assayed this at a high hematocrit level (2.5%) **(Figure 3C)**. A significant (fourfold) loss in supernatant activity was observed for the DHA-derived supernatant harvested at 6 h. Unlike DHA, for which a loss in activity was already observed after as little as 6 h, the hybrids lost their activity at 24 h. For hybrid **1**, a threefold loss in activity was observed after 24 h, while for hybrid **2** a tenfold loss in activity was observed after 24 h (**Figure 3D**). In a separate experiment, we determined that both CQ-sensitive and CQ-resistant parasites had similar susceptibility to the drug-derived supernatants using a pLDH readout.

To test the association between the antimalarial activity of hybrid **1** supernatants and chemical stability, the concentration of hybrid **1** in these supernatants was quantified by HPLC and fluorescence in a microplate reader. The concentration of hybrid **1** remained constant under incubation with PBS over the analyzed time (**Figure 3E**, Figure S1). In the presence of uRBC at hematocrit 2.5%, the concentration of hybrid **1** remained the same for over 6 h, while at 24 h it decreased from 500 to 256 nM. Moreover, we defined the slopes calculated from linear regression fits to accurately reflect the effective decay in drug concentration, because peroxides could covalently bind to proteins present in the supernatant. With this metric, the concentration of hybrid **1** was estimated to have decreased between 0.16 h and 24 h of incubation. Maximum rates of reduction of between 56.4% and 65.4% were determined by HPLC and fluorescence microplate reader, respectively (Figure S1). This decrease in the concentration of hybrid **1** is comparable to the shift in IC_50_ values of hybrid **1** in the supernatants. These conditions confirmed that the cell-host-mediated degradation of the hybrids was slower than that of DHA.

Next, we used the *in vitro* metabolism assessed in liver microsomes to gain insights into the hepatic degradation of the hybrids. Assays were performed in the presence of mouse, rat, and human microsomes, and artesunate was used as a representative ART drug. **Table 1** shows the percent of drug remaining after 30 min, the calculated intrinsic clearance (Cl_int_), and degradation half-time. We observed that artesunate remained >90% intact in mouse, rat, and human liver microsomes. Moreover, artesunate showed a low clearance rate. In the literature, artesunate is categorized as a drug of low to intermediate hepatic clearance [49, 50]. Among the compounds, artesunate showed the lowest clearance (Cl_int_ of 11.6 µL/min/mg of protein) and hybrid **2** the highest (Cl_int_ between 186.88 and 422.86 µL/min/mg of protein). Hybrid **1** remained over 70% intact in the mouse and rat liver microsomes, while in the human liver microsomes, hybrid **1** was metabolized more than artesunate. The rate of metabolic degradation for hybrid **1** was faster in human liver than in mouse or rat liver microsomes, possibly because of marked differences in cytochrome P450 expression and catalytic activity among distinct species.

**Table 1:**
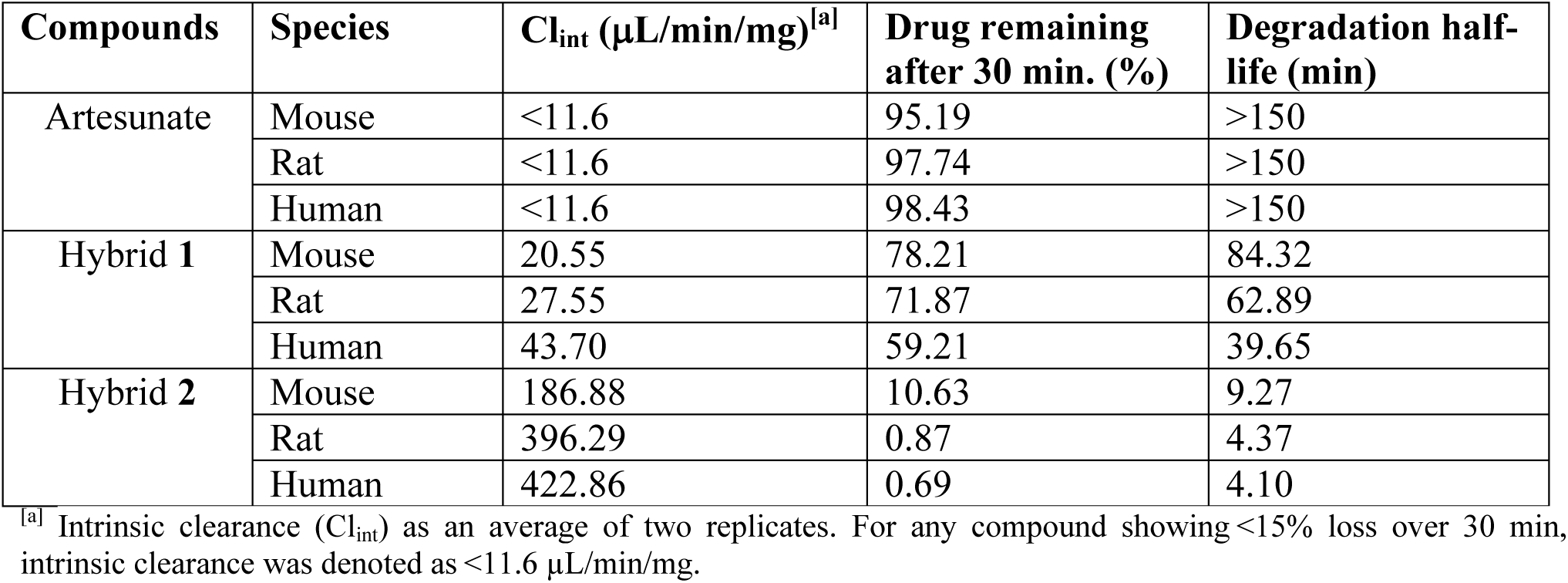
*In vitro* metabolism of drugs in the presence of mouse, rat, and human liver microsomes at physiological pH.

| Compounds | Species | $Cl_{int}$ ($\mu\text{L}/\text{min}/\text{mg}$)<sup>[a]</sup> | Drug remaining after 30 min. (%) | Degradation half-life (min) |
| --- | --- | --- | --- | --- |
| Artesunate | Mouse | <11.6 | 95.19 | >150 |
|  | Rat | <11.6 | 97.74 | >150 |
|  | Human | <11.6 | 98.43 | >150 |
| Hybrid **1** | Mouse | 20.55 | 78.21 | 84.32 |
|  | Rat | 27.55 | 71.87 | 62.89 |
|  | Human | 43.70 | 59.21 | 39.65 |
| Hybrid **2** | Mouse | 186.88 | 10.63 | 9.27 |
|  | Rat | 396.29 | 0.87 | 4.37 |
|  | Human | 422.86 | 0.69 | 4.10 |

<sup>[a]</sup> Intrinsic clearance ( $Cl_{int}$ ) as an average of two replicates. For any compound showing <15% loss over 30 min, intrinsic clearance was denoted as <11.6 $\mu\text{L}/\text{min}/\text{mg}$ .

According to the correlation of hepatic degradation with *in vivo* efficacy in *P. berghei-* infected mice (**Figure 1**, **Table 1**), hybrid **1** presented lower stability to resist hepatic metabolism than artesunate, but this occurred without compromising its biopharmaceutical properties in terms of efficacy; in fact, hybrid **1** is more efficacious than ARTs [34]. Hybrid **2** presented much lower stability to resist hepatic metabolism than artesunate, with only 10.63% drug remaining after mouse hepatic degradation. This hepatic degradation for hybrid **2** correlates well with its *in vivo* efficacy in *P. berghei-*infected mice, which is lower when given by oral administration, where extensive hepatic metabolism can take place, than by intraperitoneal injection [32].

### 3.5 Hybrids present antiplasmodial activity against ART-resistant parasites

We first established the susceptibility of ART-sensitive (F32-TEM) and ART-resistant (F32-ART) parasite strains to treatment with hybrids and DHA by determining the IC_50_ values for 48 h of continuous drug incubation. We observed that, like DHA, both molecules had IC_50_ values below 5 nM in the standard chemosensitivity assay, whatever *P. falciparum* strain was tested *in vitro* (**Table S3**). No difference was observed between the ART-resistant strain and the ART-sensitive strain. This result was expected since the mechanism of resistance to ARTs is mediated by a quiescence phenomenon (parasite cell cycle arrest during artemisinin exposure). This test based on parasite proliferation does not serve to discriminate artemisinin-sensitive parasites from resistant ones [5].

To determine whether ART resistance could impair the activity of the hybrids, a recrudescence assay was conducted [5, 10], comparing the ability of ART-resistant and ART-sensitive parasites to recover after a single pharmacologically relevant 48 h-drug exposure. Consistently, after the end of DHA exposure at 700 nM, F32-ART recovered faster than F32-TEM, with an 8-day delay being observed between the strains (**Figure 4A**). We performed the same assay for hybrids **1** and **2** and then monitored parasite growth over 30 days. As the hybrids are both artemisinin derivatives, they were tested at the same dose as DHA. Analysis of parasite recovery after the end of drug exposure revealed a delay in recrudescence time between F32-ART and F32-TEM (**Figure 4A**), demonstrating that artemisinin resistance impaired the activity of hybrids **1** and **2** and suggesting cross-resistance with ART. These data are in accordance with previously obtained results that highlight cross-resistance between ARTs and molecules harboring an endoperoxide group [22, 51, 52].

Given that hybrid **1** was more effective than DHA in delaying the rate of parasite recrudescence, we studied its drug activity in the ring-stage survival assay (RSA^0-3h^). We exposed highly synchronous ring-stage parasites (0-3 h post-invasion) harboring or not the ART-resistant phenotype and genotype to short-pulse treatment for 6 h, determining parasite survival rates after 66 h. As expected, the survival rate of the F32-TEM parasites was close to 0% after treatment with any of the three molecules (**Figure 4B**), confirming the sensitivity of this strain and the ability of hybrids **1** and **2** to act in less than 6 hours, like DHA. In contrast, the results showed that approximately 10% of the ring-stage ART-resistant parasites survived a 6 h pulse of 700 nM DHA, compared to 2% and 15% that survived a pulse with the same concentration of hybrids **1** and **2**, respectively (**Figure 4B**). These data show that parasites harboring resistance to ART are more susceptible to treatment with hybrid **1** than to treatment with DHA or hybrid **2**.

We then examined whether hybrids **1** and **2** can kill non-proliferating parasites in a quiescent state induced by ARTs. In the quiescent-stage survival assay (QSA), parasite quiescence was first induced by DHA treatment, before adding the compound to be tested (still in the presence of DHA to maintain quiescence), and then recrudescence was carefully monitored (**Figure 4C)**. Parasites treated by DHA alone can typically reach initial parasitemia between days 8 and 9 after the end of treatment, with a further delay of 6 days in recrudescence in the presence of a tested compound being the threshold of effectiveness to kill quiescent parasites [11]. Hybrid **1** seemed to delay parasite recrudescence more than hybrid **2**, which is consistent with its superior activity against proliferating parasites in the RSA^0-3h^ and reduced susceptibility to cause cross-resistance with ART. However, the delays of recrudescence observed for the two hybrids are close (0.5 and 3 days’ delay for hybrids **1** and **2**, respectively) and less the 6-day threshold, which would indicate no activity against ART-resistant parasites in a quiescent state (**Figure 4D)**. As a positive drug control, atovaquone, known to be active on quiescent parasites, is routinely tested in the QSA and has shown a delay of recrudescence of up to 10 days (data not shown) [11].

### 3.6 Hybrids perturb heme redox homeostasis and detoxification into hemozoin

Given that the hybrids are slowly degraded by RBCs and this affects their antiplasmodial activity, we tested the likelihood that this might alter their effectiveness in inhibiting heme detoxification into hemozoin and subsequently perturbing heme redox homeostasis compared to the parental drug DHA. β-hematin inhibitory activity (BHIA) was assayed to quantify both mechanisms (**Figure 5A**). Peroxide-containing drugs can present BHIA in heme-initiated assays, where a reduction of the peroxide bridge by heme produces hematin-drug adducts, but not in hematin-initiated assays [17, 18, 43]. Hematin-drug adducts inhibit the formation of β-hematin crystals by stacking hematin. Concomitant to this inhibition, the formation of these adducts is encompassed by the oxidation of iron(II) to iron(III). As a result of an augmentation in soluble heme species, which are redox-active, an imbalance in heme redox homeostasis takes place. Therefore, these BHIA assays can specifically interrogate any drug perturbation in heme detoxification into hemozoin crystals and in heme redox homeostasis.

**Figure 5:**
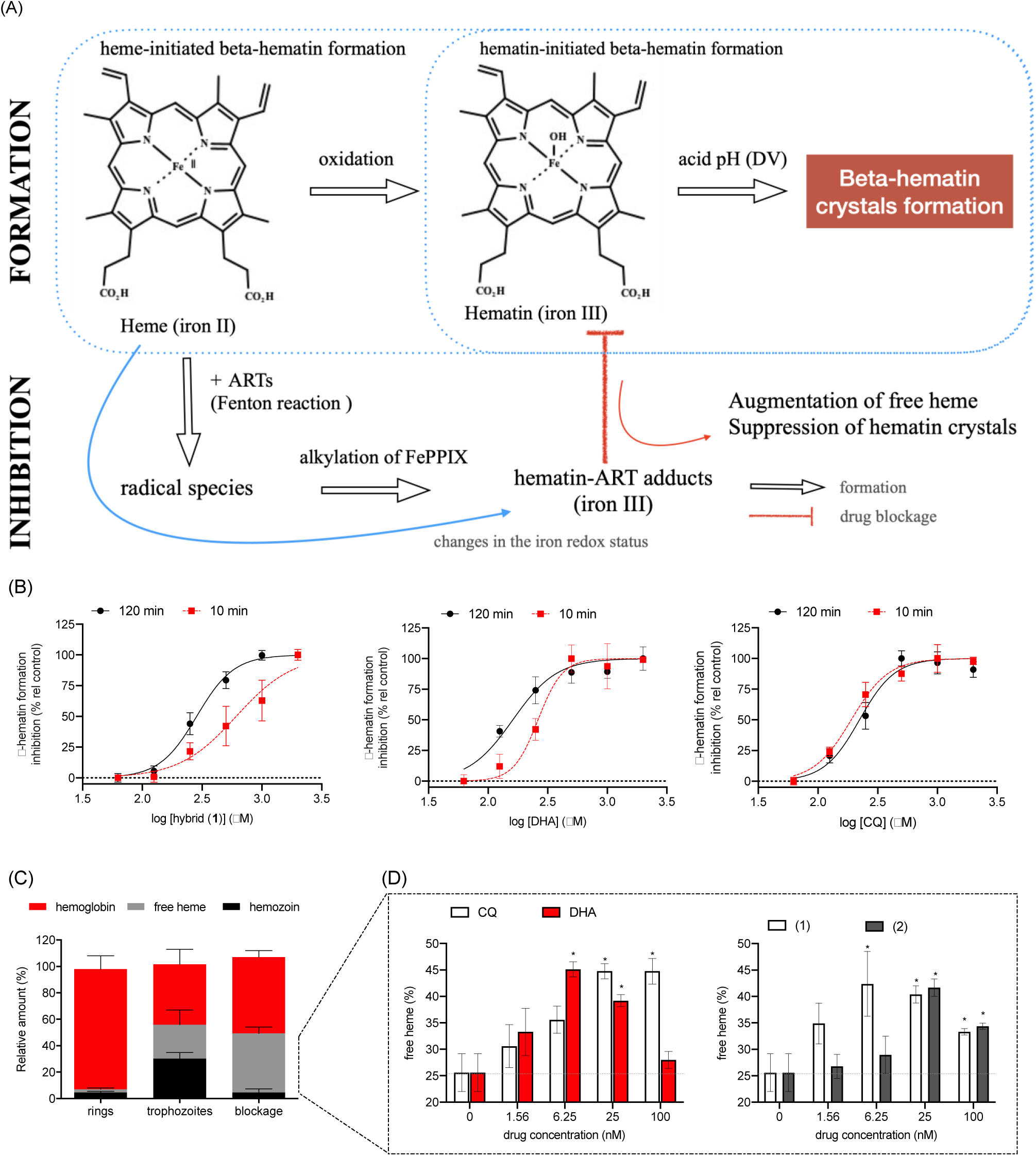
Disruption of heme detoxification and homeostasis by DHA and hybrids. Panel (A) illustrates the β-hematin inhibitory activity (BHIA) assays. Panel (B) shows that DHA and hybrid **1** have a potent BHIA in the heme-initiated β-hematin formation assay (heme is generated *in situ* in the presence of GSH as a reductant). Significant time-dependence of inhibitory activity is observed for hybrid **1**, but less for DHA and none for CQ. Values are the mean and error bars are the standard deviation from two independent experiments, each one using three replicates. Panel (C) shows the phenotype of heme detoxification in the 3D7 strain of *P. falciparum* either treated or untreated with CQ at 25 nM. Panel (D) shows the drug-concentration effect on the level of free heme. Treatment was performed at ring stages and incubation lasted until late-trophozoite stages (32 h post-infection) before the cell lysate fractionation assay. The percentage of free heme is relative to the sum of hemoglobin, free heme, and hemozoin. Values are the means and error bars are the standard deviation from two independent experiments, each one using two technical replicates. \**p* <0.05 *versus* untreated (two-way ANOVA and Bonferroni post-test). CQ = chloroquine. DHA = dihydroartemisinin. FePPIX = iron protoporphyrin IX. **Figures S6** and **S7** (supporting material) show the detailed results.

DHA and hybrid **1** did not show any BHIA in the hematin-initiated assay, while CQ showed potent activity (Figure S6). In contrast, we observed that in the heme-initiated assay, strong BHIA was observed for DHA and hybrid **1 (Figure 5B)**, with IC_50_ values derived from BHIA revealing that DHA and hybrid **1** are equipotent. Next, we examined whether the heme-induced reduction of the peroxide bridge was slower in hybrid **1** than in DHA. In comparison to the standard 120-minute incubation time for the reductive step (i.e., heme plus drugs without acid buffer), incubation as short as 10 minutes decreased the potency of BHIA threefold for hybrid **1**, slightly for DHA, but not for CQ (control). We interpreted the strong BHIA observed for DHA and hybrid **1** as being the consequence of peroxide reduction by heme and the formation of the respective hematin-drug adducts, since only the adducts can potently inhibit *β*-hematin crystals. A shorter incubation time of 10 minutes for the reductive step may reduce the abundance of hematin-drug adduct formation for hybrid **1** and subsequently less inhibition of β-hematin.

A delay in the formation of hematin-drug adducts may compromise the effectiveness of a peroxide-containing drug to suppress heme detoxification into hemozoin crystals. To interrogate this, we exposed ring-stage ART-sensitive *P. falciparum* parasites to various drug concentrations and determined the levels of free heme, hemoglobin, and hemozoin. In this cell lysate fractionation assay, the ratio of these heme-derived species changed following the progression of the parasite life cycle from ring stages into trophozoites. Compared to untreated parasites, drug blockage of heme detoxification was specifically detected by a significant increase in the levels of free heme **(Figure 5 C)**. Both CQ and DHA treatment increased free heme levels in a concentration-dependent manner **(Figure 5D)**. Under treatment with the hybrid drugs, increased free heme levels were observed, with hybrid (**2**) being less effective than hybrid (**1**) **(Figure 5D)**. Comparatively, both DHA and hybrid (**1**) increased free heme levels, with a peak of effectiveness at drug concentrations ranging from 6.2 to 25 nM, indicating that DHA and hybrid **1** are almost equipotent. Therefore, the previously determined dissimilarity in cell-host-mediated degradation between both hybrids **1** and **2** and DHA does not interfere with the drugs’ ability to inhibit heme detoxification into hemozoin.

Antimalarial 4-aminoquinolines and peroxide-containing drugs have been shown to inhibit heme detoxification in BHIA assays with similar potency [32, 43]. In the cell lysate fractionation assays, we noticed that in all tested concentrations of CQ, an increase in free heme levels was concomitant to a decrease in hemozoin content [29, 53]. However, for peroxide-containing drugs, this same profile was observed in drug concentrations up to 25 nM, while in a concentration of 100 nM, we observed increased hemoglobin levels. Differences in parasite growth inhibition according to cell potency (IC_50_ of 14.5 nM for CQ *versus* 5.0 nM for DHA) might affect the drugs’ capacity to affect heme detoxification into hemozoin. Moreover, differences in drug concentrations might shift the outcomes of heme detoxification. Here, we observed that at 100 nM, DHA increased hemoglobin levels by presumably inhibiting hemoglobin catabolism (Figure S7). Similarly, a recent study has observed that DHA can inhibit parasite growth by a mechanism of ferroptosis in concentrations up to 200 nM but not in higher concentrations that reflect the plasmatic concentration of DHA in patients [54].

## 4 DISCUSSION

ARTs are powerful antimalarial drugs bioactivated by heme, which is enriched in the *plasmodium* environment. They are degraded into radical species capable of alkylating mainly heme [16, 19], as well as a variety of molecules [32]. However, parasites can evade ART-based treatment by arresting their growth in ring stages and subsequently escaping the damage of the short-lived drug-induced radical species. Here, we have demonstrated that the peroxide stability and, consequently, the cell-host-mediated degradation of ART-based hybrid drugs **1** and **2** exceeds that of DHA, resulting in an improvement of potency against resistant parasites.

Previously, hybrids **1** and **2** were identified as having *in vitro* and *in vivo* potency superior to their pharmacophoric units of CQ, ART, and artesunate, and to display potency comparable to DHA, as a drug screening threshold [29, 32, 34]. To understand the reasons for the potency enhancement, we investigated whether the hybrids share the same phenotype of antimalarial activity as ARTs. As expected, the speed of activity against the parasites was relatively fast for the hybrids and DHA in comparison to the slow-acting drug atovaquone. Moreover, the hybrids and DHA possess stage-specific activity, killing ring-stage parasites. This phenotype of activity is broadly consistent with the bioavailability of heme in the early stages of the asexual blood stage life cycle—a requirement for activating ARTs and causing extensive alkylation and cell damage resulting in a fast reduction of parasite viability [47, 55].

We hypothesized that the rate of homolysis of the peroxide bond in the hybrids might be different from that of the parental ARTs. Using multiple experiments, we delineated cell-host-mediated degradation as a critical dissimilarity between hybrids **1** and **2** and the parental ARTs. We observed that cell-host-mediated drug degradation is consistently observed for all endoperoxide drugs but not for the 4-aminoquinoline drug AQ. Drug degradation was found to be dependent on the time of exposure and bioavailability of heme, with the highest degradation being found at the high levels of heme and for the longest exposure times, which is not entirely surprising. However, the observation of reduced cell-host-mediated degradation for the hybrids *versus* DHA is surprising, since both are endoperoxide-containing structures and are derived from the 1,2,4-trioxane present in ARTs.

Given that the hybrids have a lower cell-host-mediated degradation rate, we further examined whether the steps of peroxide activation by heme might proceed differently for the hybrids compared to DHA. IC_50_ values derived from BHIA revealed that the potency of hybrid **1** in alkylating heme species responsible for inhibiting the formation of β-hematin crystals is similar to that of DHA. Besides heme, other pathways may contribute to peroxide degradation, such as metabolism in plasma, pH in the cytosol, and hepatic metabolism [12–15]. While we cannot formally exclude all degradation pathways here, we identified that stability towards hepatic metabolism, assessed through microsome assays, did not differ significantly between artesunate and hybrid **1**. This leads us to the assumption that heme-mediated activation and degradation of hybrids is the most important pathway. A subtle increase in the IC_50_ value at a shorter drug incubation time in the BHIA assays was observed for hybrid **1**, implying that peroxide activation by heme may proceed slower for hybrid **1** than for parental DHA.

Clearly, the kinetics of heme-mediated degradation may not be the only key feature in controlling the outcome of antimalarial activity; the emerging degradation products may have an impact too [56]. A well-known product of ART degradation is hematin-drug adducts [16, 57, 58], which in recent years have been shown to effectively kill ART-resistant parasites in the RSA^0-3h^ [17, 18]. Formation of hematin-hybrid adducts from hybrid **1** and **2** was inferred from our BHIA assays. Moreover, we observed in parasite cells that hybrids can inhibit heme detoxification with a potency comparable to or higher than DHA. Any endogenous hematin-drug adducts produced in the parasite cells likely inhibit heme detoxification [17, 18]. Based on these findings, we suggest that hematin-hybrid adducts may be just as important for antiparasitic activity as the hematin-DHA adduct. Currently, the exact structures and relative abundances of hematin-hybrid adducts remain to be defined.

Multiple studies have shown that ozonides (1,2,4-trioxolanes) are more stable in overcoming cell-host-mediated degradation than ARTs (1,2,4-trioxanes) [23, 36], but this is less clear for 1,2,4,5-tetraoxane-derived drugs [21, 59]. A computational study has shown that the stability of the peroxide bond is greater for ARTs and tetraoxanes than it is for ozonides [60]. Thus, it is plausible that the increased stability of ozonides in cell-host-mediated degradation is due to their improved drug-like properties and pharmacokinetics in comparison with those of ARTs. In fact, it has been proposed that a peroxide-based drug of improved stability in cell-host-mediated degradation could confer these drugs the ability to efficiently kill ART-resistant parasites in RSA^0-3h^ [23, 36]. Our study provides important data on both points by directly showing that hybrid **1** has greater chemical stability of the peroxide bridge and can efficiently kill ART-resistant parasites in RSA^0-3h^. Hybrid **1** showed a significantly slower parasite-mediated degradation rate than hybrid **2**, the latter being degraded to a lesser degree than DHA; however, hybrid (**2**) was not effective at killing ART-resistant parasites in RSA^0-3h^. This lack of effectiveness could be because of significant degradation and therefore loss of activity against sensitive parasites at high heme levels (namely, activity depending on hematocrit). In contrast, hybrid **1** was degraded to a significantly lesser degree and only a subtle loss in activity was observed (**Figure 6**). Our findings are consistent with two recent reports showing that a chemical functionalization of the 1,2,4-trioxane warhead in ARTs could lend hybrids improved stability and antimalarial activity [33, 35].

**Figure 6:**
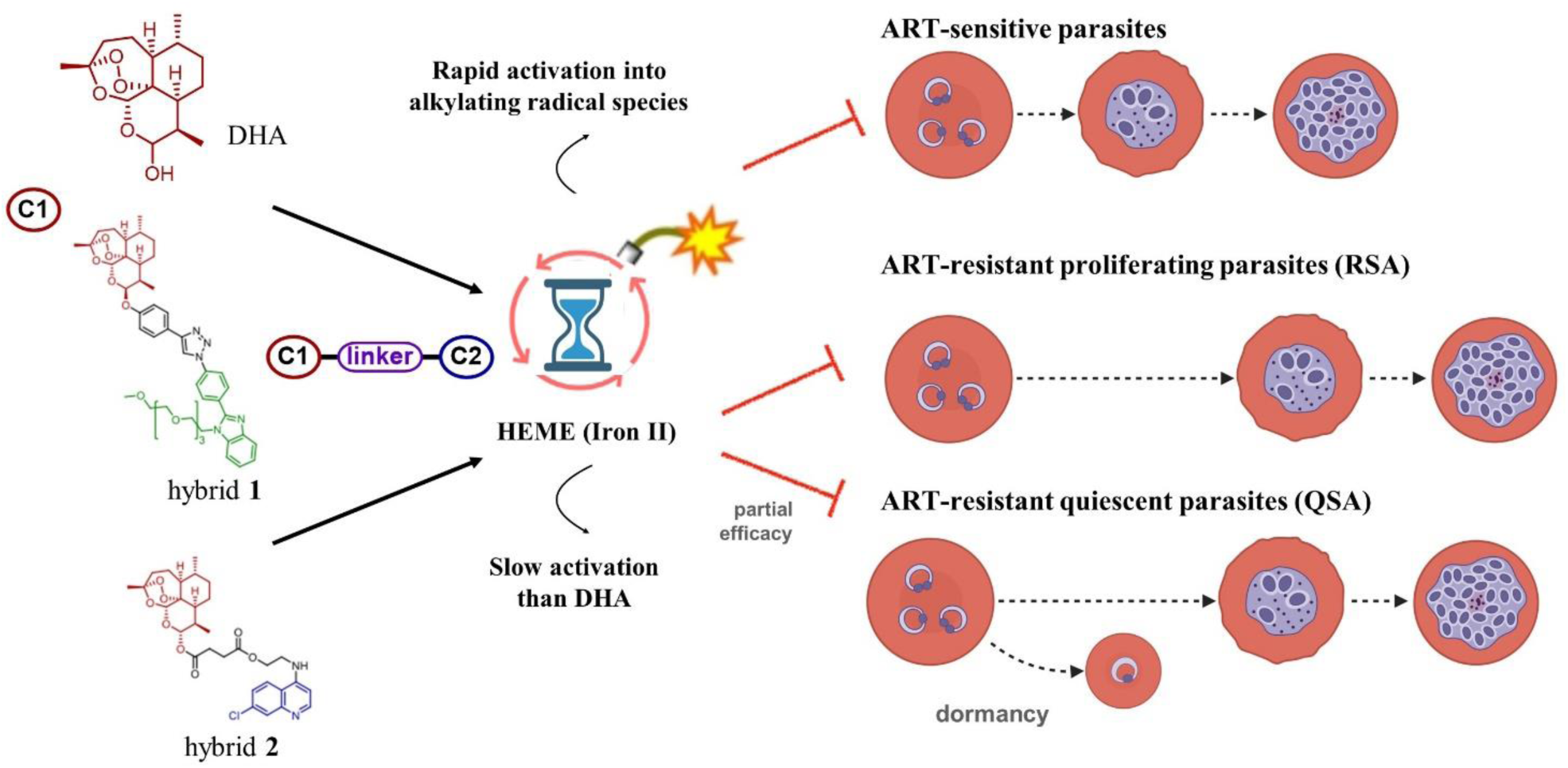
Proposed action of antimalarial peroxide-containing drugs against different parasite phenotypes of *Plasmodium*. A peroxide bond is activated by heme (iron II) through a Fenton reaction, which produces radical species that alkylate a variety of molecules and thereby toxify the parasite cells. One consequence of this drug activation is to efficiently kill ART-sensitive parasites, as exemplified by DHA and hybrids **1** and **2**. However, ART-resistant parasites show a delayed clearance phenotype, with the ring stages arresting growth upon exposure to ART. If slowly activated, peroxide drugs such as hybrid **1** could potentially kill ART-resistant parasites in the RSA, when parasites are in a proliferative state at the time of drug exposure. However, when parasites are at the quiescent stage, they are only partially affected by hybrid **1** and are unaffected by hybrid **2**.

The hepatic metabolism rate of hybrid **1** was similar to that of artesunate, but much higher in hybrid **2**. Hybrid **1** is linked via an aromatic triazole unit that is not prone to hydrolysis or enzymatic cleavage, while hybrid **2** is linked via a cleavable ester moiety. The linker stability correlates well with *in vivo* efficacy in infected mice, as hybrid **1** is more potent than hybrid **2** and ART [29, 32, 34]. While hepatic metabolism and host-mediated degradation could potentially be fine-tuned by chemical modifications and molecular hybridization, as demonstrated here, we recognize that any increase in chemical functionalization and lipophilicity may change other biopharmaceutical parameters [25, 26, 58]. For instance, we observed that our and other designed ART-based hybrids may require a high dosage for effective treatment in mice [32–35].

Ideally, any new peroxide-containing drug should be capable of being combined to maximize drug interactions with ARTs in ART-based combination therapy (ACT). Regarding this, we have shown that hybrids **1** and **2** both display a summation of activity when employed in combination with drug partners CQ and MFQ. This profile has also been reported for DHA, ART, and artesunate, as well as for long-acting ozonides [43, 56], suggesting that any peroxide drug of slow host-mediated degradation can reproduce the capability of ARTs in drug combination. However, the unresolved question is the emergence of parasites in the quiescent state after treatment with the ACT regimen [10, 11]. Our screens revealed that hybrid **1** displays strong activity against proliferating parasites (in RSA^0-3h^) but no activity against quiescent parasites (in QSA). Conversely, trioxane/quinoline hybrid **2** did not show any activity against proliferating parasites in the RSA^0-3h^. A previous study has observed low effectiveness of trioxane/quinoline hybrids (with a similar phenotype of activity and structure to hybrid **2)** in RSA^0-3h^, but no direct measurements of cell-host-mediated degradation were available in this work for comparison [51]. Currently, there are few antimalarial drugs that are efficient against quiescent parasites. Interestingly, some studies have found 4-aminoquinoline drugs to have weak activity against quiescent parasites [11]. Much work is needed to broaden the scope of peroxide drugs that kill quiescent parasites.

## 5. CONCLUSIONS

To mitigate the spread of ART resistance, there is an urgent need to make improvements to current ACT regimens. We demonstrated two hybrids with fast-acting antiplasmodial action that efficiently kill young parasites by affecting the mechanisms of heme detoxification and heme redox homeostasis. Importantly, the hybrids reproduced the most important phenotypic characteristics of ARTs, whether given as monotherapy or in combination with other drugs. Notably, we showed that the hybrids undergo classic cell-host-mediated degradation, which is intrinsically dependent on the levels of bioavailable heme, and that hybrid drug **1** overcomes fast degradation and presents higher activity against ART-resistant parasites than DHA. Our multiple approaches consistently suggest that the key feature of ozonides and tetraoxanes—slower cell-host-mediated degradation than ARTs—was also consistently reproduced by the ART-based hybrid drug **1**. Our findings broadly confirm that the manipulation of semi-synthetic trioxanes is a valuable approach in antimalarial drug design.

## Data availability

All data are available in the main text of the manuscript and supporting information section.

## Author contributions

Study design, S.B.T., F.B.V., and D.R.M.M.; experimental work, H.C.Q., L.H., J.M., L.P., M.C.B.S., G.A.D., and S.D.-A.; writing—original draft preparation, H.C.Q., L.H., and D.R.M.M.; writing—review and editing, L.P., F.B.V., N.B., K.C., S.B.T., and D.R.M.M.; funding acquisition, F.B.V., N.B., K.C., S.B.T., and D.R.M.M. All authors have revised and approved the final version of manuscript.

## Informed consent statement

Blood donor consent was waived since erythrocytes were used for *P. falciparum* culture and no personal data were collected.

## Acknowledgments

S.D.-A. and N.B. thank the Immunohematology and Transfusion Medicine Service, Department of Laboratory Medicine, and ASST Grande Ospedale Metropolitano Niguarda (Milan) for providing red blood cells for the parasite culture. H.C.Q. and D.R.M.M. thank Fiocruz for the flow cytometry data acquisition. We further wish to thank Anne Robert and Jean-Michel Augereau from CNRS au Laboratoire de Chimie de Coordination (LCC) de Toulouse (France) for their helpful inputs and revision of the manuscript.

## Supporting information

**Figure S1:**
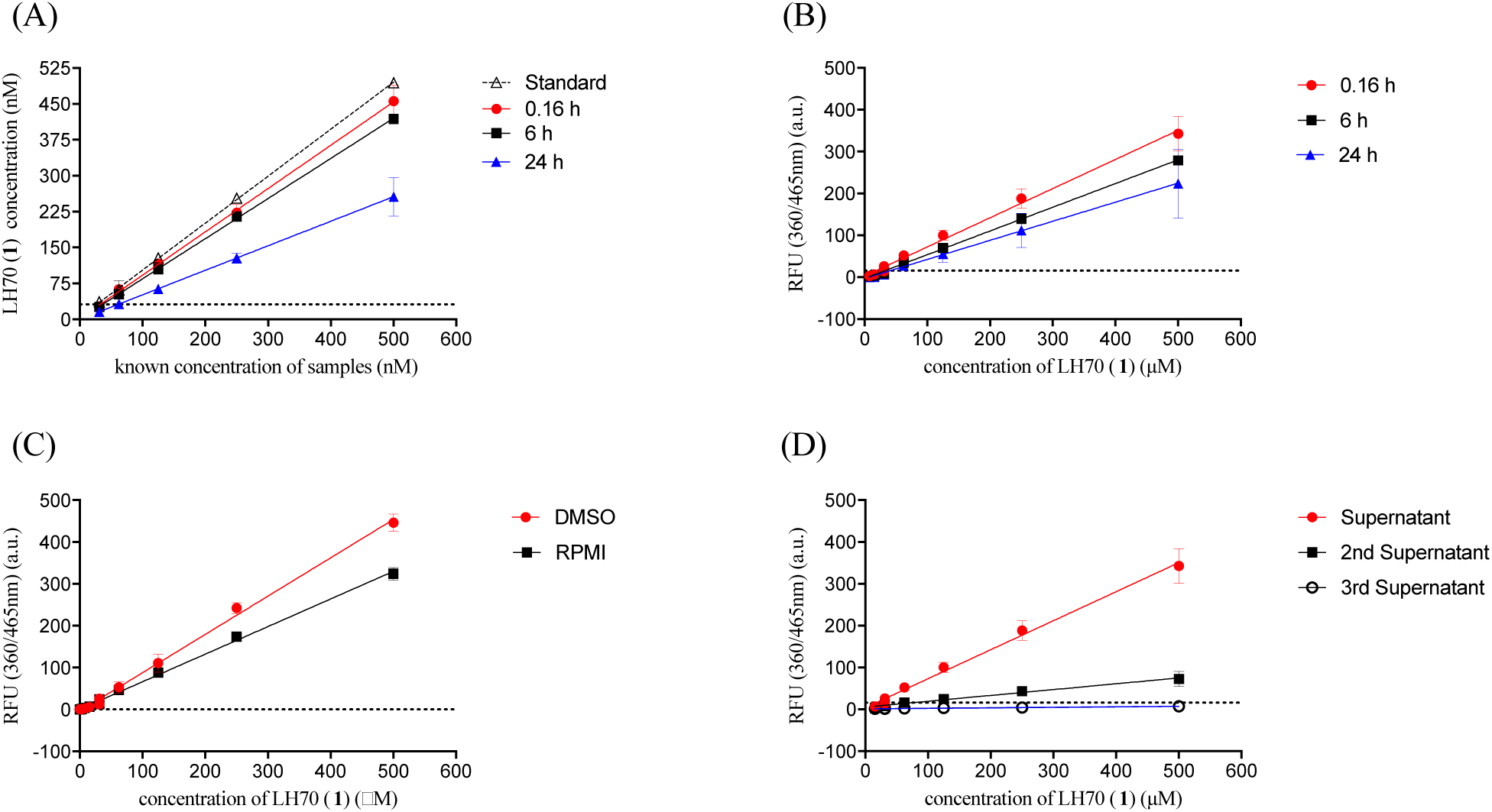
Quantification of hybrid LH70 (**1**) by fluorescence and HPLC in the uRBC-derived supernatants. Panel (A) shows the quantification of hybrid LH70 (**1**) by HPLC (Shimadzu, C18 column and UV-Vis detector set at 215 nm) in the uRBC-derived supernatants harvested at indicated incubation times. Standard represents a sample of compound freshly dissolved in DMSO and then diluted in RPMI. Panel (B) shows the quantification of hybrid LH70 (**1**) by fluorescent signal measured in a plate reader (Fluoroskan, Thermo) in the uRBC-derived supernatants harvested at indicated incubation times. Panel (C) shows the quantification of hybrid LH70 (**1**) by fluorescence of the compound freshly dissolved in pure DMSO or dissolved in DMSO and then diluted in RPMI (final concentration of DMSO 0.5% v/v). Panel (D) shows the quantification of hybrid LH70 (**1**) using a fluorescence plate reader. The drug was quantified in the supernatant of the cell culture after the second and third steps of medium replacement and washing out. In all panels, values are mean and error bars are standard deviation of one experiment performed in two replicates; black dotted line means the limit of detection. RFU = relative fluorescence units.

**Table S1:** Stage-specificity of antiparasitic activity in the asexual blood stages assayed against the CQ-sensitive strain NF54 of *P. falciparum*. Data associated to Figure 2A in the main text of the manuscript.

| Compounds | NF54 strain of *P. falciparum* (IC<sub>50</sub>, nM, median ± SEM) |  |  |  |
| --- | --- | --- | --- | --- |
|  | Ring stages <sup>[a]</sup> | Asynchronous <sup>[b]</sup> | 72 h <sup>[c]</sup> | *p* values <sup>[d]</sup> |
| LH70 (**1**) | 1.5 ± 0.3 | 3.5 ± 0.3 | 3.0 ± 0.3 | - |
| 163A (**2**) | 9.0 ± 1.5 | 19.1 ± 1.8 | 8.1 ± 1.0 | <0.05 |
| DHA | 5.0 ± 0.5 | 4.6 ± 0.9 | 2.1 ± 0.6 | - |
| MFQ | 37.2 ± 5.2 | 61.6 ± 11.9 | 20.2 ± 1.3 | <0.05 |

<sup>[a]</sup> Synchronized ring-stage parasites were incubated in the presence of compounds for 6 h, drugs were removed by washing, and plates were incubated for an additional 66 h. <sup>[b]</sup> Asynchronous parasites were incubated in the presence of compounds for 6 h, drugs were removed by washing, and plates were incubated for an additional 66 h. <sup>[c]</sup> Asynchronous parasites were incubated in the presence of compounds for 72 h. <sup>[a,b]</sup> Experiments were performed in parallel. <sup>[d]</sup> Values (rings *versus* asynchronous) were significantly different as determined by Student's *t* test. Parasite viability was determined by parasite lactate dehydrogenase. Data are from three experiments performed in duplicate. MFQ = mefloquine; DHA = dihydroartemisinin; SEM = standard error of the mean.

**Figure S2:**
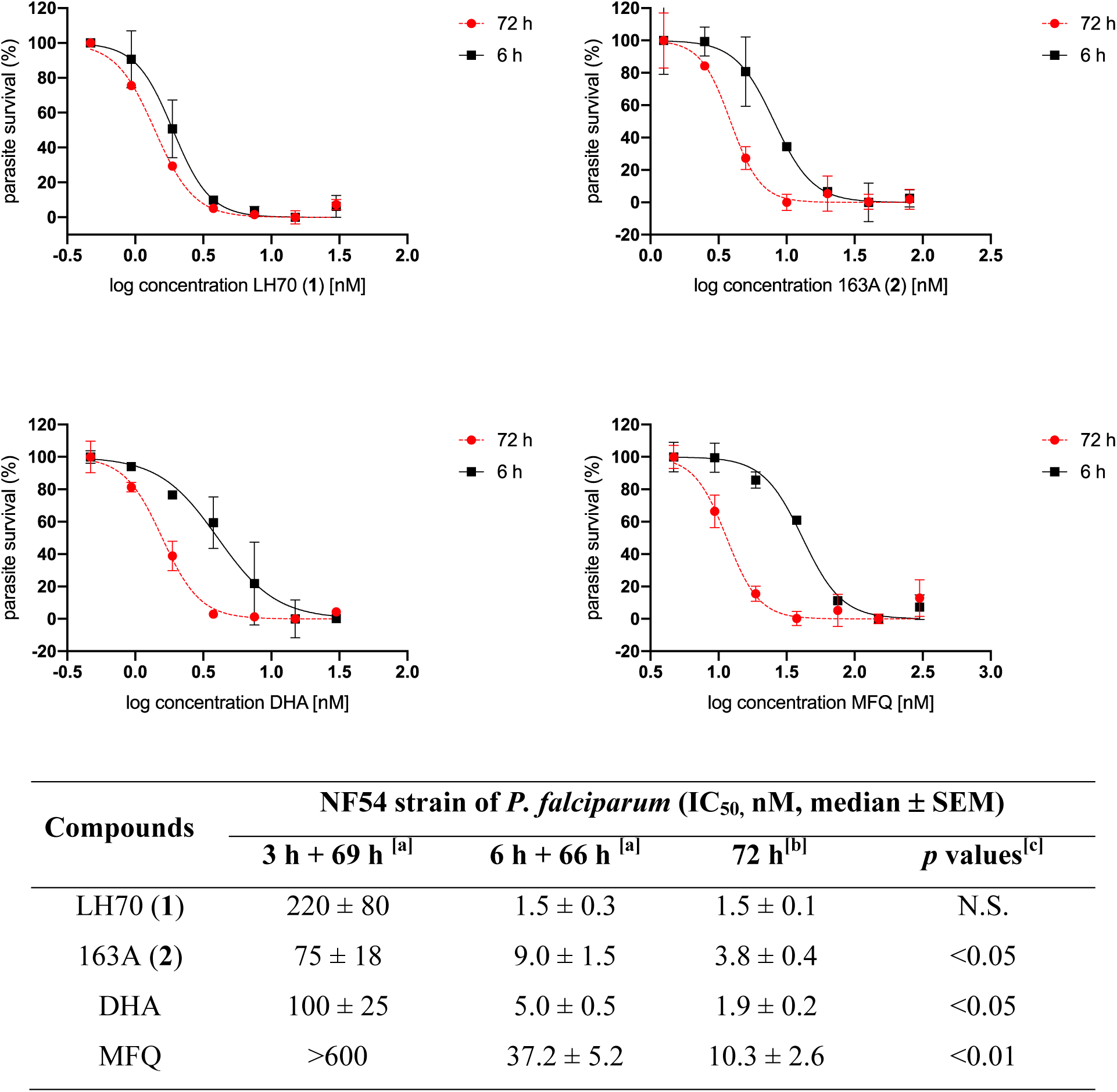
Exposure time dependence of antiparasitic activity against ring stages of CQ-sensitive strain NF54 of *P. falciparum*. Data associated to Figure 2B in the main text of the manuscript. Footnotes for table: ^[a]^ Synchronized ring-stage parasites were incubated in the presence of compounds at indicated times, drugs were removed by washing, and plates were incubated until 72 h.^[b]^ Parasites were incubated in the presence of compounds for 72 h. ^[a,b]^ Experiments were performed in parallel (paired). Parasite viability was determined by parasite lactate dehydrogenase (pLDH) assay. Data are from three experiments performed in duplicate. ^[c]^ Values (6 h versus 72 h) were significantly different as determined by Student’s *t* test. MFQ = mefloquine. DHA = dihydroartemisinin; SEM = standard error of the mean; N.S. = not statistically significant.

**Figure S3:**
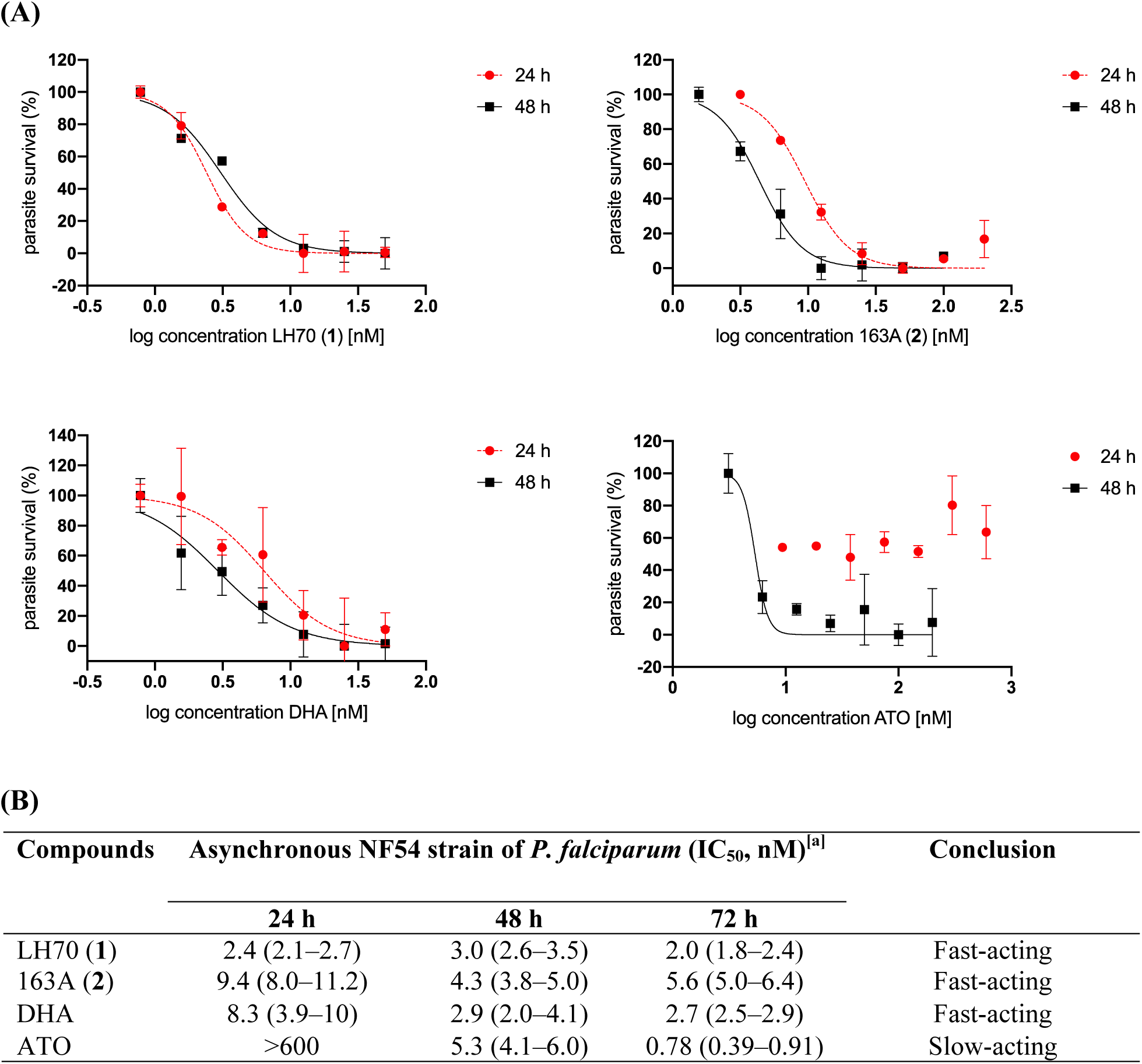
Speed of antiparasitic activity of hybrids (**1**) and (**2**) against CQ-sensitive NF54 strain of *P. falciparum*. Panel A: Representative response-concentration curves. Panel B: A table summarizing the results. Footnotes for table: ^[a]^ Parasite viability was accessed at each indicated time (24, 48, or 72 h) after addition of drugs. Values are the mean and 95% confidence interval of one experiment using each concentration of compounds in duplicate. Parasite viability was determined by parasite lactate dehydrogenase (pLDH) assay. DHA = dihydroartemisinin; ATO = atovaquone.

**Figure S4.**
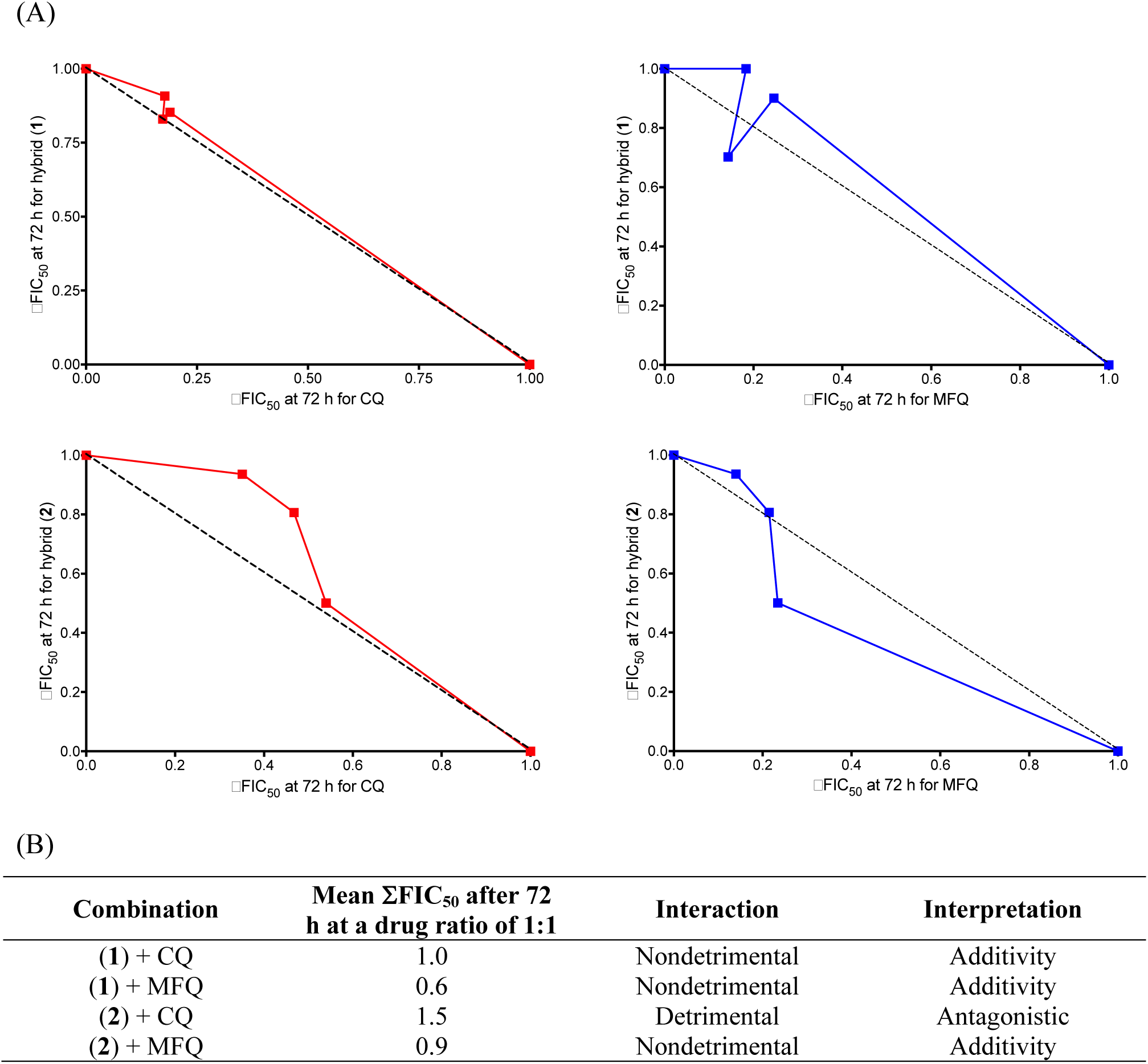
*In vitro* interactions of CQ and MFQ plus hybrids (1) and (2) against the NF54 strain of *P. falciparum*. Panel A shows representative isobologram plots. Values represent one experiment; each concentration of drug combination was tested in technical duplicate. Black dashed line indicates **Σ**FIC = 1 (absolute additivity). Panel B shows a table summarizing the **Σ**FIC indices derived from panel A. Values are the means of two independent experiments. **Σ**FIC = fractional inhibitory concentration. FIC index <1 is synergistic, ∼1 is additive, and >1 is antagonistic. Nondetrimental interactions are 0.25 <**Σ**FIC <1.25. CQ = chloroquine; MFQ = mefloquine.

**Table S2.** IC_50_ of drugs alone or in combination with CQ or MFQ. Data associated to Figure S4.

| Ring stages of NF54 strain *P. falciparum* (IC<sub>50</sub>, nM)<sup>[a]</sup> |  |  |  |  |
| --- | --- | --- | --- | --- |
| Drugs alone |  | Drugs in combination at different ratios |  |  |
|  |  | 3:1 | 1:1 | 1:3 |
| Hybrid (1) | CQ | (1):CQ | (1):CQ | (1):CQ |
| 2.2 (2.0–2.4) | 10.5 (7.5–14.8) | 1.9 (1.6–2.1) | 1.8 (1.5–2.2) | 2.0 (1.7–2.3) |
| Hybrid (1) | MFQ | (1):MFQ | (1):MFQ | (1):MFQ |
| 2.0 (1.9–2.2) | 10.3 (8.6–12.3) | 1.9 (1.7–2.0) | 1.5 (1.3–1.7) | 2.5 (2.0–3.1) |
| Hybrid (2) | CQ | (2):CQ | (2):CQ | (2):CQ |
| 4.7 (4.6–4.8) | 10.4 (9.2–11.7) | 4.9 (4.5–5.2) | 4.8 (4.4–5.1) | 4.4 (3.7–5.1) |
| Hybrid (2) | MFQ | (2):MFQ | (2):MFQ | (2):MFQ |
| 4.8 (4.5–5.1) | 17.0 (14.0–20.6) | 4.0 (3.8–4.3) | 3.4 (2.8–4.1) | 2.5 (2.0–2.9) |

<sup>[a]</sup> Values are the mean and 95% confidence interval of one experiment for each concentration of compounds in duplicate. CQ = chloroquine; MFQ = mefloquine.

**Figure S5:**
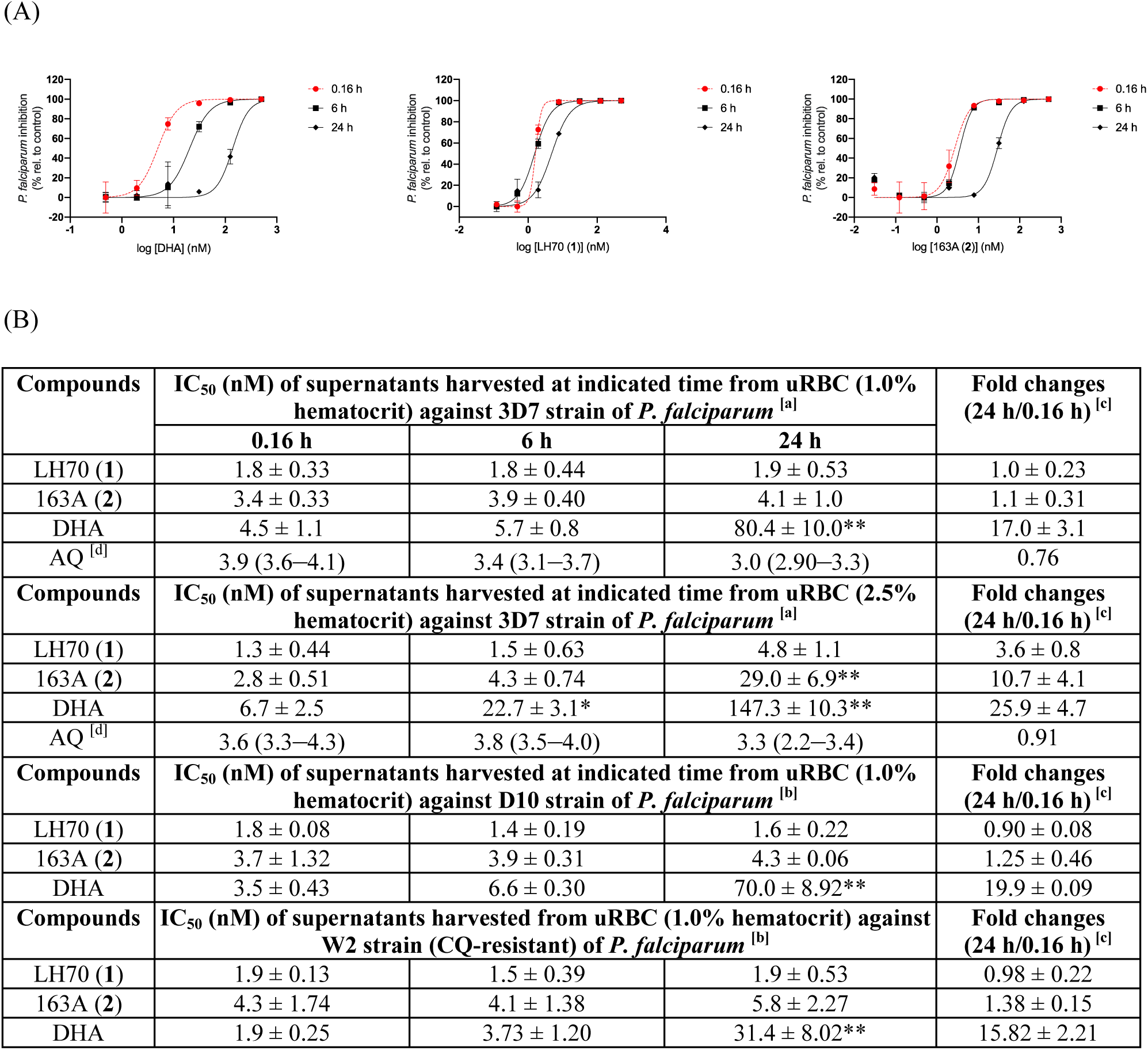
Data associated to Figure 3 in the main text of the manuscript. Panel A: Representative response-concentration curves of the antiplasmodial activity of supernatant-derived uRBC (2.5% hematocrit) harvested at indicated times (0.16, 6, and 24 h). Drug activity on parasite viability was determined 72 h after parasite incubation (3D7 strain, asynchronous). Values were subtracted from control (without treatment), transformed, and normalized to calculate nonlinear fit. Parasite viability was determined by SYBR Green I. Values are mean and error bars are the standard deviation of a single experiment, each concentration in duplicate. Panel B: A table summarizing the IC_50_ values. Footnotes: ^[a]^ Parasite viability was determined by SYBR Green I. ^[b]^ Parasite viability was determined by pLDH readout. Values are median ± SEM of three independent experiments, each concentration in duplicate. ^[c]^ Calculated as the ratio of IC_50_ at 24 h to IC_50_ at 0.16 h.^[d]^ These values are expressed as mean and 95% confidence interval (CI) of one experiment, each concentration in duplicate. AQ = amodiaquine; DHA = dihydroartemisinin; uRBC = uninfected red blood cells. Unpaired two-tailed *t*-test at 95% CI: \**p* <0.05; \*\**p* <0.01.

**Table S3:**
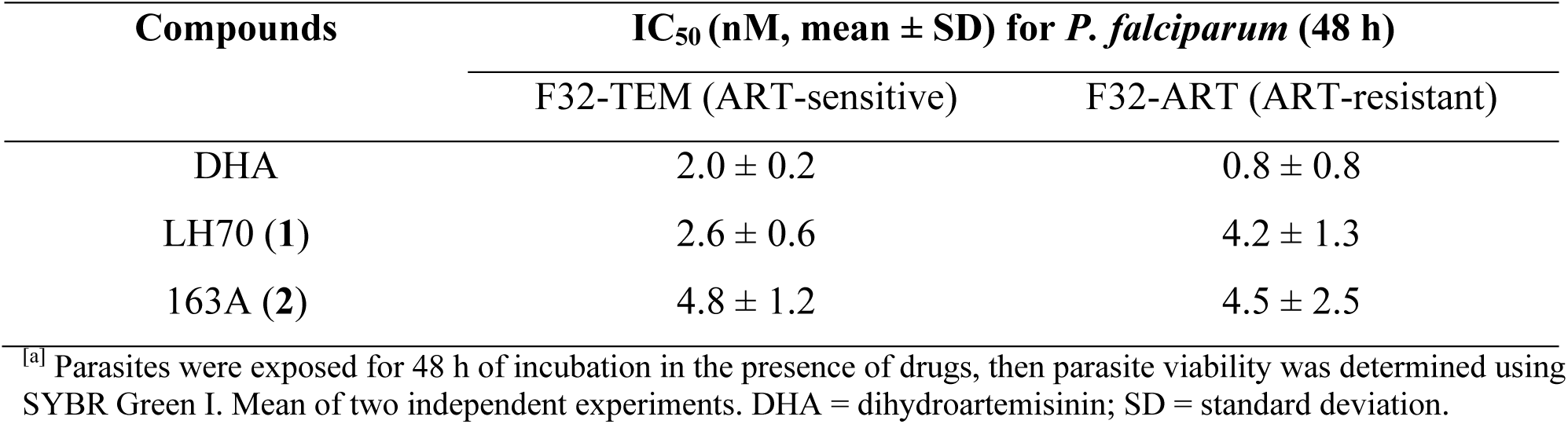
Summary of IC_50_ values against F32-ART strain (ART-resistant) and its isogenic laboratory control F32-TEM strain (ART-sensitive) of *P. falciparum*.

**Figure S6:**
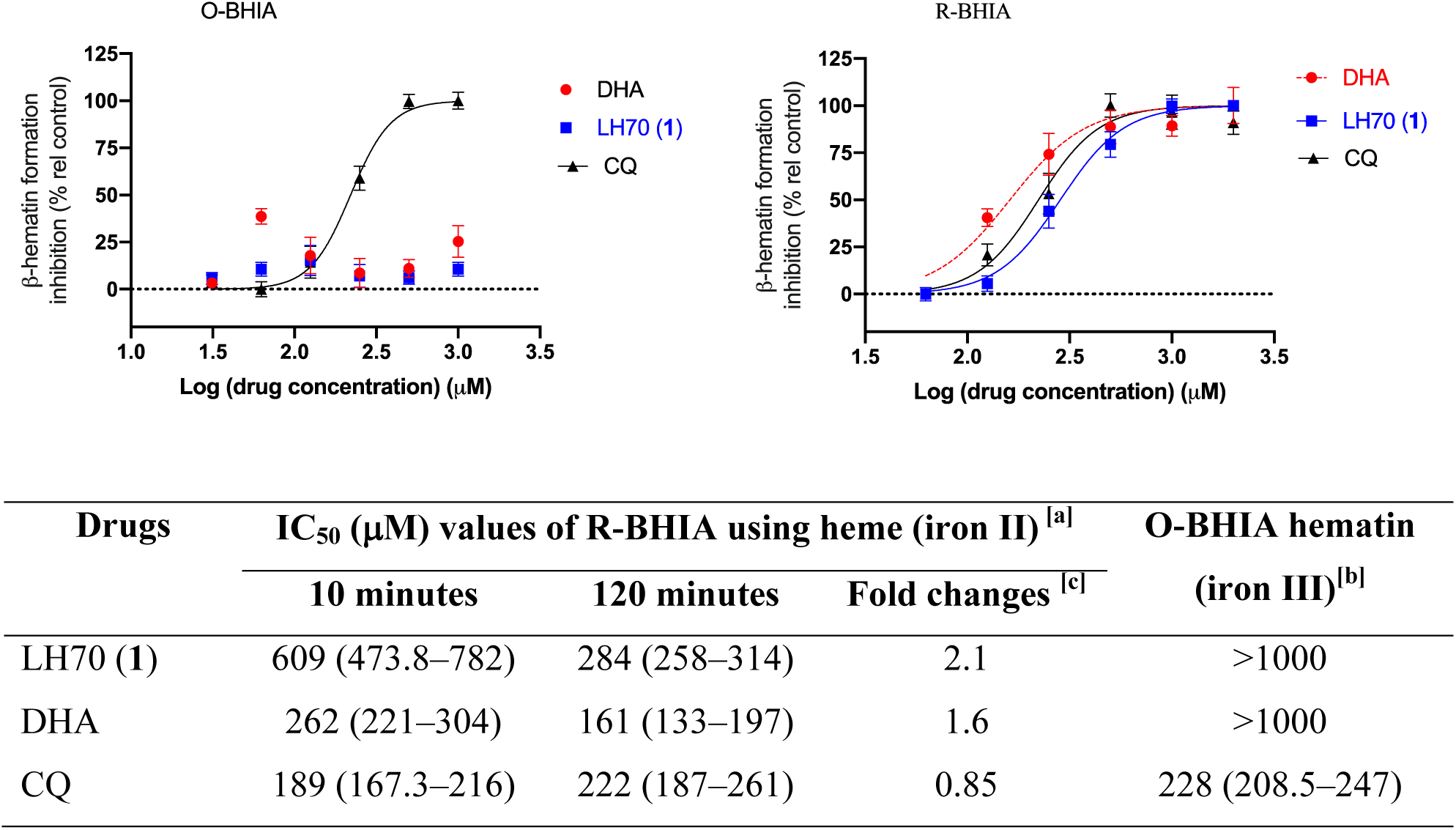
A summary of IC_50_ values derived from β-hematin inhibitory activity (BHIA). Footnotes for table: ^[a,b]^ Values are mean and 95% confidence interval of one experiment, each concentration in triplicate. ^[a]^ Using heme (Fe[II]PPIX[Cl]) as a starting reagent. ^[b]^ Using hematin (Fe[III]PPIX[OH]) as a starting reagent. ^[c]^ Ratio of IC_50_ (10 minutes) to IC_50_ (120 minutes). CQ = chloroquine. R-BHIA = reducing β-hematin inhibitory activity. O-BHIA = oxidizing β-hematin inhibitory activity.

**Figure S7:**
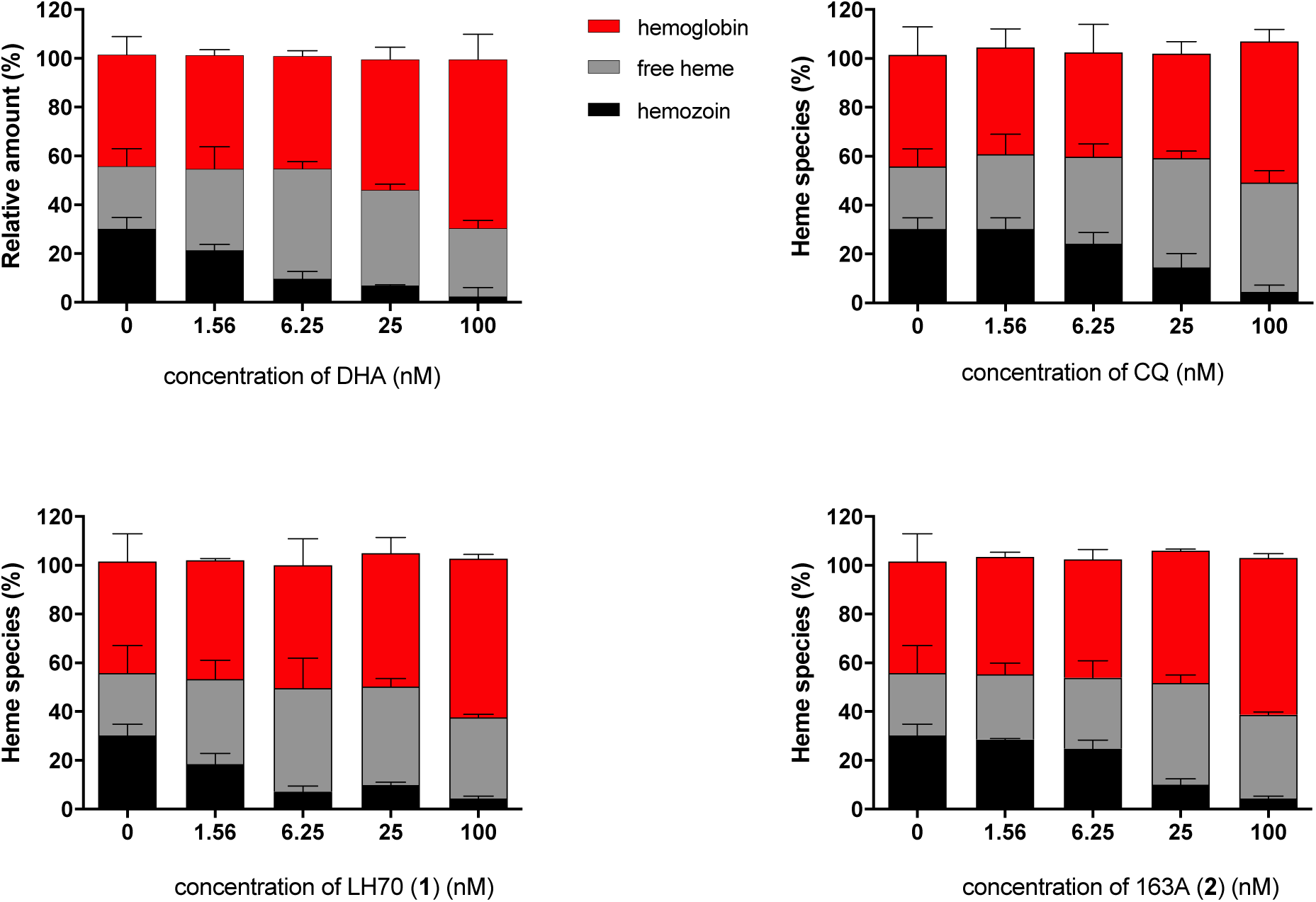
Quantification of heme species involved in the heme detoxification of *P. falciparum* by treatment with DHA, CQ, and hybrids. Values are the mean and error bars are the standard deviation from two independent experiments, each one using two biological replicates. CQ = chloroquine. DHA = dihydroartemisinin. Data associated to Figure 5 in the main text of the manuscript.

## References

[1] Wang J, Xu C, Lun ZR, Meshnick SR. Unpacking ‘Artemisinin Resistance’. Trends Pharmacol Sci. 2017 Jun;38(6):506–511. doi: 10.1016/j.tips.2017.03.007.

[2] Ma N, Zhang Z, Liao F, Jiang T, Tu Y. The birth of artemisinin. Pharmacol Ther. 2020 Dec;216:107658. doi: 10.1016/j.pharmthera.2020.107658.

[3] Do Dondorp AM, Nosten F, Yi P, Das D, Phyo AP, Tarning J, Lwin KM, Ariey F, Hanpithakpong W, Lee SJ, Ringwald P, Silamut K, Imwong M, Chotivanich K, Lim P, Herdman T, An SS, Yeung S, Singhasivanon P, Day NP, Lindegardh N, Socheat D, White NJ. Artemisinin resistance in Plasmodium falciparum malaria. N Engl J Med. 2009 Jul 30;361(5):455–67. doi: 10.1056/NEJMoa0808859. Erratum in: N Engl J Med. 2009 Oct 22;361(17):1714. PMID: 19641202; PMCID: PMC3495232.

[4] Mok S, Ashley EA, Ferreira PE, Zhu L, Lin Z, Yeo T, Chotivanich K, Imwong M, Pukrittayakamee S, Dhorda M, Nguon C, Lim P, Amaratunga C, Suon S, Hien TT, Htut Y, Faiz MA, Onyamboko MA, Mayxay M, Newton PN, Tripura R, Woodrow CJ, Miotto O, Kwiatkowski DP, Nosten F, Day NP, Preiser PR, White NJ, Dondorp AM, Fairhurst RM, Bozdech Z. Drug resistance. Population transcriptomics of human malaria parasites reveals the mechanism of artemisinin resistance. Science. 2015 Jan 23;347(6220):431–5. doi: 10.1126/science.1260403.

[5] Witkowski B, Lelièvre J, Barragán MJ, Laurent V, Su XZ, Berry A, Benoit-Vical F. Increased tolerance to artemisinin in Plasmodium falciparum is mediated by a quiescence mechanism. Antimicrob Agents Chemother. 2010 May;54(5):1872–7. doi: 10.1128/AAC.01636-09.

[6] Ariey F, Witkowski B, Amaratunga C, Beghain J, Langlois AC, Khim N, Kim S, Duru V, Bouchier C, Ma L, Lim P, Leang R, Duong S, Sreng S, Suon S, Chuor CM, Bout DM, Ménard S, Rogers WO, Genton B, Fandeur T, Miotto O, Ringwald P, Le Bras J, Berry A, Barale JC, Fairhurst RM, Benoit-Vical F, Mercereau-Puijalon O, Ménard D. A molecular marker of artemisinin-resistant Plasmodium falciparum malaria. Nature. 2014 Jan 2;505(7481):50–5. doi: 10.1038/nature12876.

[7] Pacheco MA, Kadakia ER, Chaudhary Z, Perkins DJ, Kelley J, Ravishankar S, Cranfield M, Talundzic E, Udhayakumar V, Escalante AA. Evolution and Genetic Diversity of the *k13* Gene Associated with Artemisinin Delayed Parasite Clearance in Plasmodium falciparum. Antimicrob Agents Chemother. 2019 Jul 25;63(8):e02550–18. doi: 10.1128/AAC.02550-18.

[8] Hassett MR, Roepe PD. Origin and Spread of Evolving Artemisinin-Resistant *Plasmodium falciparum*Malarial Parasites in Southeast Asia. Am J Trop Med Hyg. 2019 Dec;101(6):1204–1211. doi: 10.4269/ajtmh.19-0379.

[9] Mok S, Stokes BH, Gnädig NF, Ross LS, Yeo T, Amaratunga C, Allman E, Solyakov L, Bottrill AR, Tripathi J, Fairhurst RM, Llinás M, Bozdech Z, Tobin AB, Fidock DA. Artemisinin-resistant K13 mutations rewire Plasmodium falciparum’s intra-erythrocytic metabolic program to enhance survival. Nat Commun. 2021 Jan 22;12(1):530. doi: 10.1038/s41467-020-20805-w

[10] Ménard S, Ben Haddou T, Ramadani AP, Ariey F, Iriart X, Beghain J, Bouchier C, Witkowski B, Berry A, Mercereau-Puijalon O, Benoit-Vical F. Induction of Multidrug Tolerance in Plasmodium falciparum by Extended Artemisinin Pressure. Emerg Infect Dis. 2015 Oct;21(10):1733–41. doi: 10.3201/eid2110.150682

[11] Reyser T, Paloque L, Ouji M, Nguyen M, Ménard S, Witkowski B, Augereau JM, Benoit-Vical F. Identification of compounds active against quiescent artemisinin-resistant Plasmodium falciparum parasites via the quiescent-stage survival assay (QSA). J Antimicrob Chemother. 2020 Oct 1;75(10):2826–2834. doi: 10.1093/jac/dkaa250

[12] Batty KT, Gibbons PL, Davis TM, Ilett KF. Pharmacokinetics of dihydroartemisinin in a murine malaria model. Am J Trop Med Hyg. 2008 Apr;78(4):641–2.

[13] Parapini S, Olliaro P, Navaratnam V, Taramelli D, Basilico N. Stability of the antimalarial drug dihydroartemisinin under physiologically relevant conditions: implications for clinical treatment and pharmacokinetic and in vitro assays. Antimicrob Agents Chemother. 2015 Jul;59(7):4046–52. doi: 10.1128/AAC.00183-15.

[14] Sissoko A, Vásquez-Ocmín P, Maciuk A, Barbieri D, Neveu G, Rondepierre L, Grougnet R, Leproux P, Blaud M, Hammad K, Michel S, Lavazec C, Clain J, Houzé S, Duval R. A Chemically Stable Fluorescent Mimic of Dihydroartemisinin, Artemether, and Arteether with Conserved Bioactivity and Specificity Shows High Pharmacological Relevance to the Antimalarial Drugs. ACS Infect Dis. 2020 Jul 10;6(7):1532–1547. doi: 10.1021/acsinfecdis.9b00430.

[15] Bai G, Gao Y, Liu S, Shui S, Liu G. pH-dependent rearrangement determines the iron-activation and antitumor activity of artemisinins. Free Radic Biol Med. 2021 Feb 1;163:234–242. doi: 10.1016/j.freeradbiomed.2020.12.024.

[16] Robert A, Benoit-Vical F, Claparols C, Meunier B. The antimalarial drug artemisinin alkylates heme in infected mice. Proc Natl Acad Sci U S A. 2005 Sep 20;102(38):13676–80. doi: 10.1073/pnas.0500972102.

[17] Ma W, Balta VA, West R, Newlin KN, Miljanić OŠ, Sullivan DJ, Vekilov PG, Rimer JD. A second mechanism employed by artemisinins to suppress Plasmodium falciparum hinges on inhibition of hematin crystallization. J Biol Chem. 2021 Jan-Jun;296:100123. doi: 10.1074/jbc.RA120.016115.

[18] Ma W, Balta VA, Pan W, Rimer JD, Sullivan DJ, Vekilov PG. Nonclassical mechanisms to irreversibly suppress β-hematin crystal growth. Commun Biol. 2023 Jul 27;6(1):783. doi: 10.1038/s42003-023-05046-z.

[19] Gao P., Wang J., Chen J., Gu L., Wang C., Xu L., Wang J. Profiling the Antimalarial Mechanism of Artemisinin by Identifying Crucial Target Proteins. Engineering, 2023, in press. DOI: 10.1016/j.eng.2023.06.001.

[20] Batty KT, Thu LT, Davis TM, Ilett KF, Mai TX, Hung NC, Tien NP, Powell SM, Thien HV, Binh TQ, Kim NV. A pharmacokinetic and pharmacodynamic study of intravenous vs oral artesunate in uncomplicated falciparum malaria. Br J Clin Pharmacol. 1998 Feb;45(2):123–9. doi: 10.1046/j.1365-2125.1998.00655.x

[21] O’Neill PM, Amewu RK, Charman SA, Sabbani S, Gnädig NF, Straimer J, Fidock DA, Shore ER, Roberts NL, Wong MH, Hong WD, Pidathala C, Riley C, Murphy B, Aljayyoussi G, Gamo FJ, Sanz L, Rodrigues J, Cortes CG, Herreros E, Angulo-Barturén I, Jiménez-Díaz MB, Bazaga SF, Martínez-Martínez MS, Campo B, Sharma R, Ryan E, Shackleford DM, Campbell S, Smith DA, Wirjanata G, Noviyanti R, Price RN, Marfurt J, Palmer MJ, Copple IM, Mercer AE, Ruecker A, Delves MJ, Sinden RE, Siegl P, Davies J, Rochford R, Kocken CHM, Zeeman AM, Nixon GL, Biagini GA, Ward SA. A tetraoxane-based antimalarial drug candidate that overcomes PfK13-C580Y dependent artemisinin resistance. Nat Commun. 2017 May 24;8:15159. doi: 10.1038/ncomms15159.

[22] Straimer J, Gnädig NF, Stokes BH, Ehrenberger M, Crane AA, Fidock DA. *Plasmodium falciparum* K13 Mutations Differentially Impact Ozonide Susceptibility and Parasite Fitness *In Vitro*. mBio. 2017 Apr 11;8(2):e00172–17. doi: 10.1128/mBio.00172-17.

[23] Giannangelo C, Stingelin L, Yang T, Tilley L, Charman SA, Creek DJ. Parasite-Mediated Degradation of Synthetic Ozonide Antimalarials Impacts In Vitro Antimalarial Activity. Antimicrob Agents Chemother. 2018 Feb 23;62(3):e01566–17. doi: 10.1128/AAC.01566-17.

[24] Siddiqui G, Giannangelo C, De Paoli A, Schuh AK, Heimsch KC, Anderson D, Brown TG, MacRaild CA, Wu J, Wang X, Dong Y, Vennerstrom JL, Becker K, Creek DJ. Peroxide Antimalarial Drugs Target Redox Homeostasis in Plasmodium falciparum Infected Red Blood Cells. ACS Infect Dis. 2022 Jan 14;8(1):210–226. doi: 10.1021/acsinfecdis.1c00550

[25] Blank BR, Gonciarz RL, Talukder P, Gut J, Legac J, Rosenthal PJ, Renslo AR. Antimalarial Trioxolanes with Superior Drug-Like Properties and In Vivo Efficacy. ACS Infect Dis. 2020 Jul 10;6(7):1827–1835. doi: 10.1021/acsinfecdis.0c00064

[26] Blank BR, Gut J, Rosenthal PJ, Renslo AR. Artefenomel Regioisomer RLA-3107 Is a Promising Lead for the Discovery of Next-Generation Endoperoxide Antimalarials. ACS Med Chem Lett. 2023 Apr 4;14(4):493–498. doi: 10.1021/acsmedchemlett.3c00039.

[27] Benoit-Vical F, Lelièvre J, Berry A, Deymier C, Dechy-Cabaret O, Cazelles J, Loup C, Robert A, Magnaval JF, Meunier B. Trioxaquines are new antimalarial agents active on all erythrocytic forms, including gametocytes. Antimicrob Agents Chemother. 2007 Apr;51(4):1463–72. doi: 10.1128/AAC.00967-06.

[28] Coslédan F, Fraisse L, Pellet A, Guillou F, Mordmüller B, Kremsner PG, Moreno A, Mazier D, Maffrand JP, Meunier B. Selection of a trioxaquine as an antimalarial drug candidate. Proc Natl Acad Sci U S A. 2008 Nov 11;105(45):17579–84. doi: 10.1073/pnas.0804338105.

[29] Çapcı A, Lorion MM, Wang H, Simon N, Leidenberger M, Borges Silva MC, Moreira DRM, Zhu Y, Meng Y, Chen JY, Lee YM, Friedrich O, Kappes B, Wang J, Ackermann L, Tsogoeva SB. Artemisinin-(Iso)quinoline Hybrids by C-H Activation and Click Chemistry: Combating Multidrug-Resistant Malaria. Angew Chem Int Ed Engl. 2019 Sep 9;58(37):13066–13079. doi: 10.1002/anie.201907224

[30] Aratikatla EK, Kalamuddin M, Rana KC, Datta G, Asad M, Sundararaman S, Malhotra P, Mohmmed A, Bhattacharya AK. Combating multi-drug resistant malaria parasite by inhibiting falcipain-2 and heme-polymerization: Artemisinin-peptidyl vinyl phosphonate hybrid molecules as new antimalarials. Eur J Med Chem. 2021 Aug 5;220:113454. doi: 10.1016/j.ejmech.2021.113454

[31] Zhan W, Li D, Subramanyaswamy SB, Liu YJ, Yang C, Zhang H, Harris JC, Wang R, Zhu S, Rocha H, Sherman J, Qin J, Herring M, Simwela NV, Waters AP, Sukenick G, Cui L, Rodriguez A, Deng H, Nathan CF, Kirkman LA, Lin G. Dual-pharmacophore artezomibs hijack the Plasmodium ubiquitin-proteasome system to kill malaria parasites while overcoming drug resistance. Cell Chem Biol. 2023 May 18;30(5):457–469.e11. doi: 10.1016/j.chembiol.2023.04.006.

[32] Quadros, H. C., Çapcı, A., Herrmann, L., D’Alessandro, S., Fontinha, D., Azevedo, R., Villarreal, W., Basilico, N., Prudêncio, M., Tsogoeva, S. B., Moreira, D. R. M. (2021a). Studies of Potency and Efficacy of an Optimized Artemisinin-Quinoline Hybrid against Multiple Stages of the Plasmodium Life Cycle. Pharmaceuticals, 14(11), 1129. 10.3390/ph14111129

[33] Herrmann L, Leidenberger M, de Morais AS Moreira, Mai C, Capci A, Silva MCB, Plass F, Kahnt A, DRM, Kappes B, Tsogoeva SB. Autofluorescent antimalarials by hybridization of artemisinin and coumarin: in vitro/in vivo studies and live-cell imaging. Chem. Sci., 2023,14, 12941–12952. Doi: 10.1039/D3SC03661H

[34] Herrmann L, Leidenberger M, Quadros HC, Grau, BG, Jenne F, Shankara GK, Hampel F, Friedrich O, Moreira DRM, Kappes B, Tsogoeva SB. Antimalarials via Organo-Click Reaction: High In Vitro/In Vivo Activity Against Multidrug-Resistant Malaria Parasites. JACS Au, 2024, in press. Doi: 10.1021/jacsau.3c00716.

[35] Du Y, Giannangelo C, He W, Shami GJ, Zhou W, Yang T, Creek DJ, Dogovski C, Li X, Tilley L. Dimeric Artesunate Glycerophosphocholine Conjugate Nano-Assemblies as Slow-Release Antimalarials to Overcome Kelch 13 Mutant Artemisinin Resistance. Antimicrob Agents Chemother. 2022 May 17;66(5):e0206521. doi: 10.1128/aac.02065-21.

[36] Yang T, Xie SC, Cao P, Giannangelo C, McCaw J, Creek DJ, Charman SA, Klonis N, Tilley L. Comparison of the Exposure Time Dependence of the Activities of Synthetic Ozonide Antimalarials and Dihydroartemisinin against K13 Wild-Type and Mutant Plasmodium falciparum Strains. Antimicrob Agents Chemother. 2016 Jul 22;60(8):4501–10. doi: 10.1128/AAC.00574-16

[37] Makler MT, Hinrichs DJ. Measurement of the lactate dehydrogenase activity of Plasmodium falciparum as an assessment of parasitemia. Am J Trop Med Hyg. 1993 Feb;48(2):205–10. doi: 10.4269/ajtmh.1993.48.205.

[38] Lambros C, Vanderberg JP. Synchronization of Plasmodium falciparum erythrocytic stages in culture. J Parasitol. 1979 Jun;65(3):418–20.

[39] Sanz LM, Crespo B, De-Cózar C, Ding XC, Llergo JL, Burrows JN, García-Bustos JF, Gamo FJ. *P. falciparum* in vitro killing rates allow to discriminate between different antimalarial mode-of-action. PLoS One. 2012;7(2):e30949. doi: 10.1371/journal.pone.0030949.

[40] Di, L.; Kerns, E. H.; Gao, N.; Li, S. Q.; Huang, Y.; Bourassa, J. L.; Huryn, D. M. Experimental design on single-time-point high-throughput microsomal stability assay. J. Pharm. Sci. 2004, 93, 1537−1544.

[41] Witkowski B, Amaratunga C, Khim N, Sreng S, Chim P, Kim S, Lim P, Mao S, Sopha C, Sam B, Anderson JM, Duong S, Chuor CM, Taylor WR, Suon S, Mercereau-Puijalon O, Fairhurst RM, Menard D. Novel phenotypic assays for the detection of artemisinin-resistant Plasmodium falciparum malaria in Cambodia: in-vitro and ex-vivo drug-response studies. Lancet Infect Dis. 2013 Dec;13(12):1043–9. doi: 10.1016/S1473-3099(13)70252-4

[42] Combrinck JM, Fong KY, Gibhard L, Smith PJ, Wright DW, Egan TJ. Optimization of a multi-well colorimetric assay to determine haem species in Plasmodium falciparum in the presence of anti-malarials. Malar J. 2015 Jun 24;14:253. doi: 10.1186/s12936-015-0729-9.

[43] Ribbiso KA, Heller LE, Taye A, Julian E, Willems AV, Roepe PD. Artemisinin-Based Drugs Target the Plasmodium falciparum Heme Detoxification Pathway. Antimicrob Agents Chemother. 2021 Mar 18;65(4):e02137–20. doi: 10.1128/AAC.02137-20.

[44] Herrmann L, Hahn F, Grau BW, Wild M, Niesar A, Wangen C, Kataev E, Marschall M, Tsogoeva SB. Autofluorescent Artemisinin-Benzimidazole Hybrids via Organo-Click Reaction: Study of Antiviral Properties and Mode of Action in Living Cells. Chemistry. 2023 Aug 25;29(48):e202301194. doi: 10.1002/chem.202301194.

[45] Fröhlich T, Hahn F, Belmudes L, Leidenberger M, Friedrich O, Kappes B, Couté Y, Marschall M, Tsogoeva SB. Synthesis of Artemisinin-Derived Dimers, Trimers and Dendrimers: Investigation of Their Antimalarial and Antiviral Activities Including Putative Mechanisms of Action. Chemistry. 2018 Jun 7;24(32):8103–8113. doi: 10.1002/chem.201800729.

[46] Maerki S, Brun R, Charman SA, Dorn A, Matile H, Wittlin S. In vitro assessment of the pharmacodynamic properties and the partitioning of OZ277/RBx-11160 in cultures of Plasmodium falciparum. J Antimicrob Chemother. 2006 Jul;58(1):52–8. doi: 10.1093/jac/dkl209.

[47] Murithi JM, Owen ES, Istvan ES, Lee MCS, Ottilie S, Chibale K, Goldberg DE, Winzeler EA, Llinás M, Fidock DA, Vanaerschot M. Combining Stage Specificity and Metabolomic Profiling to Advance Antimalarial Drug Discovery. Cell Chem Biol. 2020 Feb 20;27(2):158–171.e3. doi: 10.1016/j.chembiol.2019.11.009.

[48] Rottmann M, Jonat B, Gumpp C, Dhingra SK, Giddins MJ, Yin X, Badolo L, Greco B, Fidock DA, Oeuvray C, Spangenberg T. Preclinical Antimalarial Combination Study of M5717, a Plasmodium falciparum Elongation Factor 2 Inhibitor, and Pyronaridine, a Hemozoin Formation Inhibitor. Antimicrob Agents Chemother. 2020 Mar 24;64(4):e02181–19. doi: 10.1128/AAC.02181-19.

[49] Li XQ, Björkman A, Andersson TB, Gustafsson LL, Masimirembwa CM. Identification of human cytochrome P(450)s that metabolise anti-parasitic drugs and predictions of in vivo drug hepatic clearance from in vitro data. Eur J Clin Pharmacol. 2003 Sep;59(5-6):429–42. doi: 10.1007/s00228-003-0636-9.

[50] Thelingwani R, Leandersson C, Bonn B, Smith P, Chibale K, Masimirembwa C. Characterisation of artemisinin-chloroquinoline hybrids for potential metabolic liabilities. Xenobiotica. 2016;46(3):234–40. doi: 10.3109/00498254.2015.1070975.

[51] Paloque L, Witkowski B, Lelièvre J, Ouji M, Ben Haddou T, Ariey F, Robert A, Augereau JM, Ménard D, Meunier B, Benoit-Vical F. Endoperoxide-based compounds: cross-resistance with artemisinins and selection of a Plasmodium falciparum lineage with a K13 non-synonymous polymorphism. J Antimicrob Chemother. 2018 Feb 1;73(2):395–403. doi: 10.1093/jac/dkx412.

[52] Ouji M, Barnoin G, Fernández Álvarez Á, Augereau JM, Hemmert C, Benoit-Vical F, Gornitzka H. Hybrid Gold(I) NHC-Artemether Complexes to Target Falciparum Malaria Parasites. Molecules. 2020 Jun 18;25(12):2817. doi: 10.3390/molecules25122817.

[53] Combrinck JM, Mabotha TE, Ncokazi KK, Ambele MA, Taylor D, Smith PJ, Hoppe HC, Egan TJ. Insights into the role of heme in the mechanism of action of antimalarials. ACS Chem Biol. 2013 Jan 18;8(1):133–7. doi: 10.1021/cb300454t.

[54] Li S, Xu W, Wang H, Tang T, Ma J, Cui Z, Shi H, Qin T, Zhou H, Li L, Jiang T, Li C. Ferroptosis plays an essential role in the antimalarial mechanism of low-dose dihydroartemisinin. Biomed Pharmacother. 2022 Apr;148:112742. doi: 10.1016/j.biopha.2022.112742.

[55] Egwu CO, Pério P, Augereau JM, Tsamesidis I, Benoit-Vical F, Reybier K. Resistance to artemisinin in falciparum malaria parasites: A redox-mediated phenomenon. Free Radic Biol Med. 2022 Feb 1;179:317–327. doi: 10.1016/j.freeradbiomed.2021.08.016

[56] Fügi MA, Wittlin S, Dong Y, Vennerstrom JL. Probing the antimalarial mechanism of artemisinin and OZ277 (arterolane) with nonperoxidic isosteres and nitroxyl radicals. Antimicrob Agents Chemother. 2010 Mar;54(3):1042–6. doi: 10.1128/AAC.01305-09.

[57] Loup C, Lelièvre J, Benoit-Vical F, Meunier B. Trioxaquines and heme-artemisinin adducts inhibit the in vitro formation of hemozoin better than chloroquine. Antimicrob Agents Chemother. 2007 Oct;51(10):3768–70. doi: 10.1128/AAC.00239-07

[58] Heller LE, Goggins E, Roepe PD. Dihydroartemisinin-Ferriprotoporphyrin IX Adduct Abundance in Plasmodium falciparum Malarial Parasites and the Relationship to Emerging Artemisinin Resistance. Biochemistry. 2018 Dec 26;57(51):6935–6945. doi: 10.1021/acs.biochem.8b00960.

[59] Woodley CM, Nixon GL, Basilico N, Parapini S, Hong WD, Ward SA, Biagini GA, Leung SC, Taramelli D, Onuma K, Hasebe T, O’Neill PM. Enantioselective Synthesis and Profiling of Potent, Nonlinear Analogues of Antimalarial Tetraoxanes E209 and N205. ACS Med Chem Lett. 2021 Jun 24;12(7):1077–1085. doi: 10.1021/acsmedchemlett.1c00031

[60] Fernández I, Robert A. Peroxide bond strength of antimalarial drugs containing an endoperoxide cycle. Relation with biological activity. Org Biomol Chem. 2011 Jun 7;9(11):4098–107. doi: 10.1039/c1ob05088e.

